# The Amsterdam Open MRI Collection, a set of multimodal MRI datasets for individual difference analyses

**DOI:** 10.1101/2020.06.16.155317

**Authors:** Lukas Snoek, Maite M. van der Miesen, Tinka Beemsterboer, Andries van der Leij, Annemarie Eigenhuis, H. Steven Scholte

## Abstract

We present the Amsterdam Open MRI Collection (AOMIC): three datasets with multimodal (3T) MRI data including structural (T1-weighted), diffusion-weighted, and (resting-state and task-based) functional BOLD MRI data, as well as detailed demographics and psychometric variables from a large set of healthy participants (*N* = 928, *N* = 226, and *N* = 216). Notably, task-based fMRI was collected during various robust paradigms (targeting naturalistic vision, emotion perception, working memory, face perception, cognitive conflict and control, and response inhibition) for which extensively annotated event-files are available. For each dataset and data modality, we provide the data in both raw and preprocessed form (both compliant with the Brain Imaging Data Structure), which were subjected to extensive (automated and manual) quality control. All data is publicly available from the Openneuro data sharing platform.

## Background & Summary

It is becoming increasingly clear that robust effects in neuroimaging studies require very large sample sizes^1,2^, especially when investigating between-subject effects^3^. With this in mind, we have run several large-scale “population imaging” MRI projects over the past decade at the University of Amsterdam, with the aim to reliably estimate the (absence) of structural and functional correlates of human behavior and mental processes. After publishing several articles using these datasets^4–7^, we believe that making the data from these projects publicly available will benefit the neuroimaging community most. To this end, we present the Amsterdam Open MRI Collection (AOMIC) — three large-scale datasets with high-quality, multimodal 3T MRI data and detailed demographic and psychometric data, which are publicly available from the Openneuro data sharing platform.

We believe that AOMIC represents a useful contribution to the growing collection of publicly available population imaging MRI datasets^8–11^. AOMIC contains a large representative dataset of the general population, “ID1000” (*N* = 928), and two large datasets with data from university students, “PIOP1” (*N* = 216) and “PIOP2” (*N = 226; P*opulation *I*maging *o*f *P*sychology). AOMIC contains MRI data from multiple modalities (structural, diffusion, and both task-based and resting-state functional MRI), concurrently measured physiological (respiratory and cardiac) data, and a variety of well-annotated demographics (age, sex, handedness, educational level, etc.), psychometric measures (intelligence, personality), and behavioral information related to the task-based fMRI runs (see Figure 1 and Table 1 for an overview). Furthermore, AOMIC offers, in addition to the raw data, also preprocessed data from well-established preprocessing and quality control pipelines, all consistently formatted according to the Brain Imaging Data Structure^12^. As such, researchers can quickly and easily prototype and implement novel secondary analyses without having to worry about quality control and preprocessing themselves.

**Figure 1.**
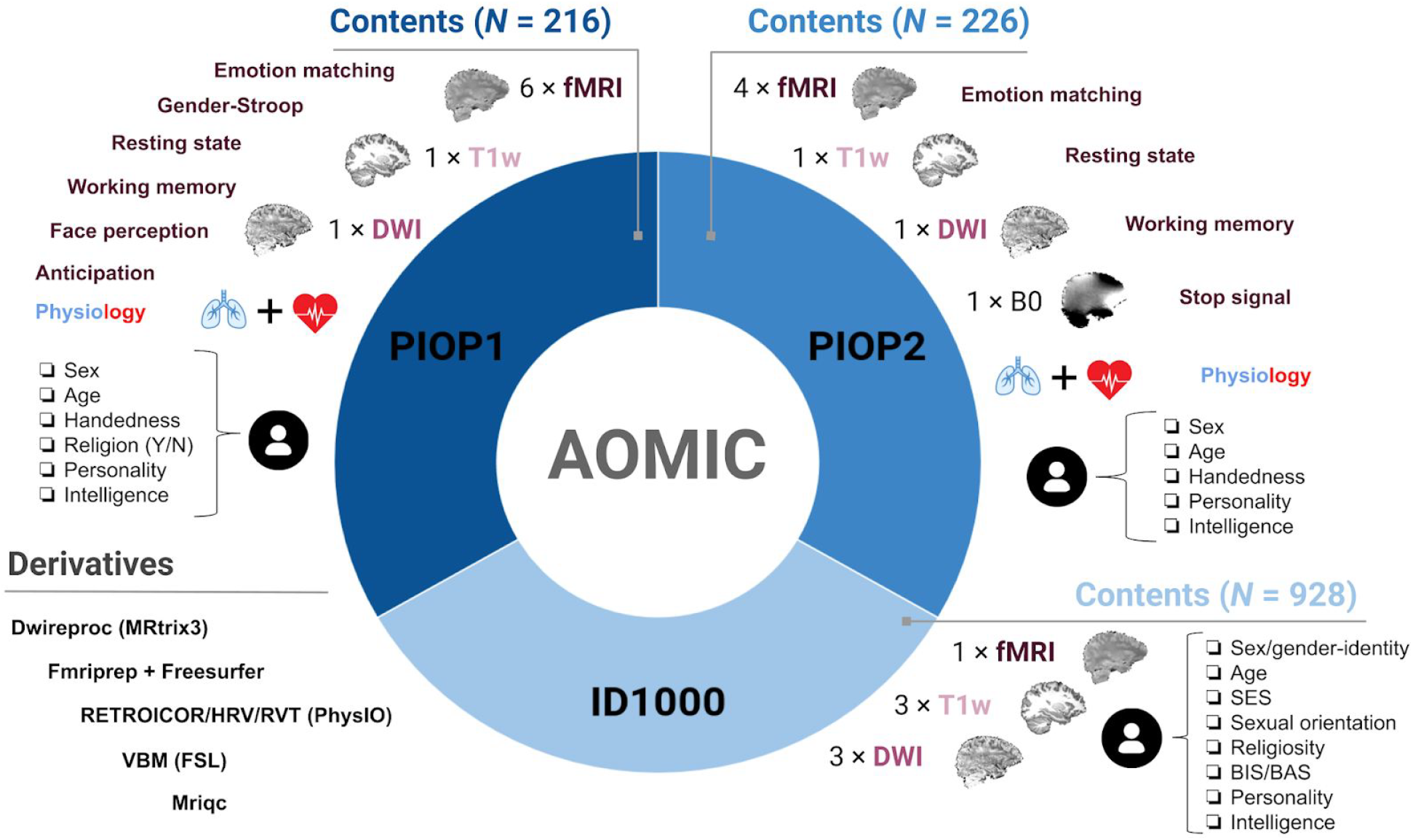
General overview of AOMIC’s contents. Each dataset (ID1000, PIOP1, PIOP2) contains multimodal MRI data, physiology (concurrent with fMRI acquisition), demographic and psychometric data, as well as a large set of “derivatives”, i.e., data derived from the original “raw” data through state-of-the-art preprocessing pipelines.

**Table 1.** Overview of the number of subjects per dataset and tasks. Mov: movie watching, RS: resting-state, Emo: emotion matching, G-str: gender-stroop, FP: face perception, WM: working memory, Antic: anticipation, Stop: stop-signal.

|  | *ID1000* |  | *PIOP1* |  |  |  |  | *PIOP2* |  |  |  |
| --- | --- | --- | --- | --- | --- | --- | --- | --- | --- | --- | --- |
| *N subj.* | 928 |  |  | 216 |  |  |  |  | 226 |  |  |
| *T1w* | 928 |  |  | 216 |  |  |  |  | 226 |  |  |
| *DWI* | 925 |  |  | 211 |  |  |  |  | 226 |  |  |
| *Fieldmap* | n/a |  |  | n/a |  |  |  |  | 226 |  |  |
|  | *Mov* | *RS* | *Emo* | *G-str* | *FP* | *WM* | *Antic* | *RS* | *Emo* | *WM* | *Stop* |
| *fMRI* | 881 | 210 | 208 | 208 | 203 | 207 | 203 | 214 | 222 | 224 | 226 |
| *Physiology* | 790 | 198 | 194 | 194 | 189 | 194 | 188 | 216 | 216 | 211 | 217 |

Due to the size and variety of the data in AOMIC, there are many ways in which it can be used for secondary analysis. One promising direction is to use the data for the development of generative and discriminative machine learning-based algorithms, which often need large datasets to train models on. Another, but related, use of AOMIC’s data is to use it as a validation dataset (rather than train-set) for already developed (machine learning) algorithms to assess the algorithm’s ability to generalize to different acquisition sites or protocols. Lastly, due to the rich set of confound variables shipped with each dataset (including physiology-derived noise regressors), AOMIC can be used to develop, test, or validate (novel) denoising methods.

## Methods

In this section, we describe the details of the data acquisition for each dataset in AOMIC. We start with a common description of the MRI scanner used to collect the data. The next two sections describe the participant characteristics, data collection protocols, and experimental paradigms (for functional MRI) separately for the ID1000 study and the PIOP studies. Then, two sections describe the recorded subject-specific variables (such as educational level, background socio-economic status, age, etc.) and psychometric measures from questionnaires and tasks (such as intelligence and personality). Finally, we describe how we standardized and preprocessed the data, yielding an extensive set of “derivatives” (i.e., data derived from the original raw data).

### Scanner details and general scanning protocol (all datasets)

Data from all three datasets were scanned on the same Philips 3T scanner (Philips, Best, the Netherlands), but underwent several upgrades in between the three studies. The ID1000 dataset was scanned on the “Intera” version, after which the scanner was upgraded to the “Achieva” version (converting a part of the signal acquisition pathway from analog to digital) on which the PIOP1 dataset was scanned. After finishing the PIOP1 study, the scanner was upgraded to the “Achieva dStream” version (with even earlier digitalization of the MR signal resulting in less noise interference), on which the PIOP2 study was scanned. All studies were scanned with a 32-channel head coil (though the head coil was upgraded at the same time as the dStream upgrade).

At the start of each scan session, a low resolution survey scan was made, which was used to determine the location of the field-of-view. For all structural (T1-weighted), fieldmap (phase-difference based B0 map), and diffusion (DWI) scans, the slice stack was not angulated. This was also the case for the functional MRI scans of ID1000 and PIOP1, but for PIOP2 the slice stack for functional MRI scans was angulated such that the eyes were excluded as much as possible in order to reduce signal dropout in orbitofrontal cortex. While the set of scans acquired for each study is relatively consistent (i.e., at least one T1-weighted anatomical scan, at least one diffusion-weighted scan, and at least one functional BOLD MRI scan), the parameters for a given scan vary slightly between the three studies. In Table 2, the parameters for the different types of scans across all three studies are listed.

**Table 2.**
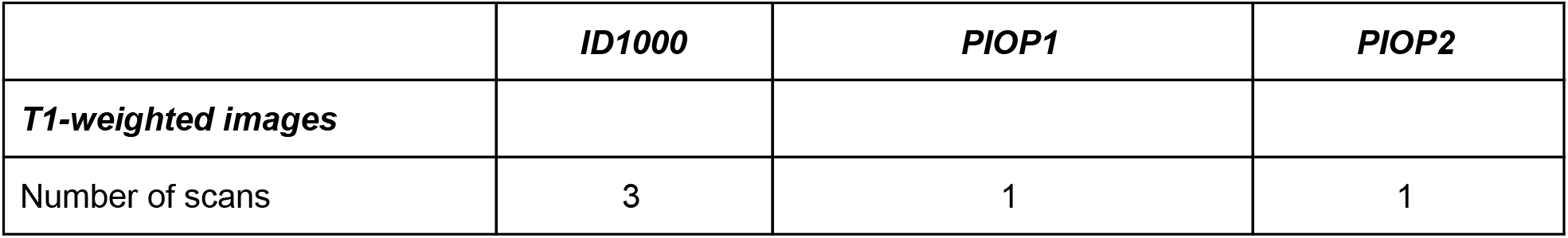
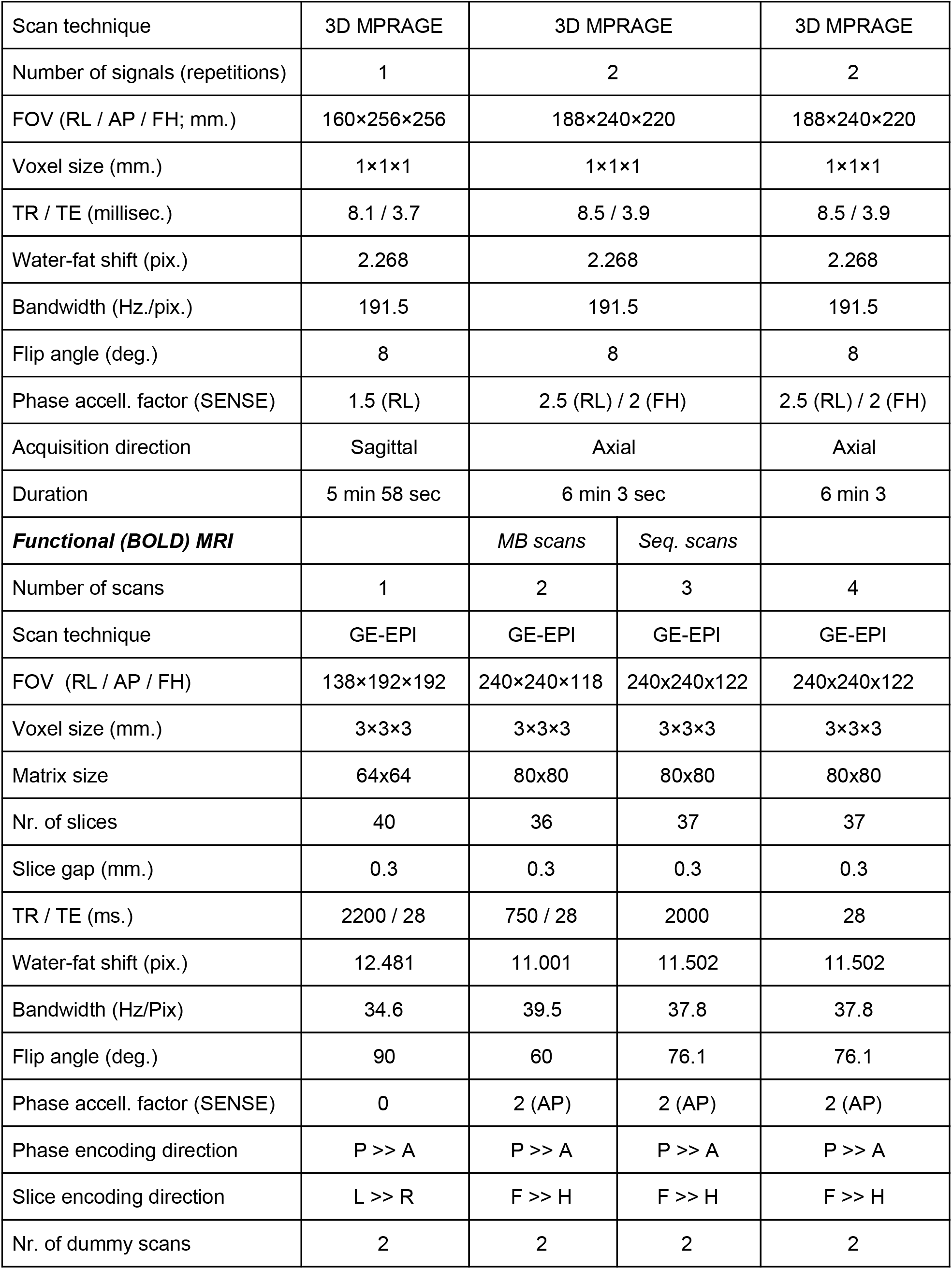
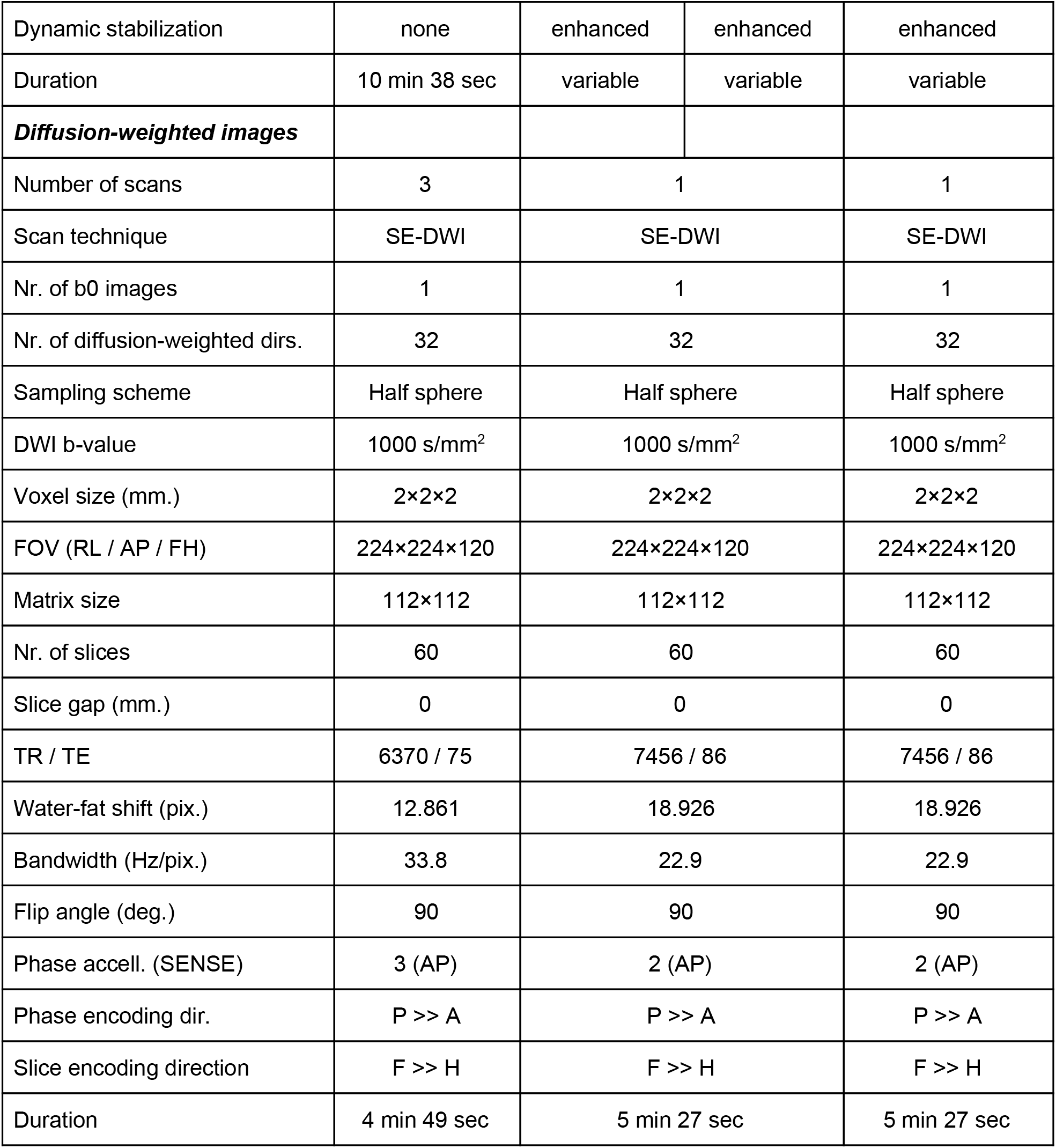
Acquisition parameters for the different scans acquired across all three datasets. MPRAGE: Magnetization Prepared Rapid Gradient Echo, FOV: field-of-view, RL: right-left, AP: anterior-posterior, FH: feet-head, GE-EPI: gradient echo-echo-planar imaging, TR: time to repetition, TE: time to echo.

|  | *ID1000* | *PIOP1* | *PIOP2* |
| --- | --- | --- | --- |
| ***T1-weighted images*** |  |  |  |
| Number of scans | 3 | 1 | 1 |

|  |  |  |  |  |
| --- | --- | --- | --- | --- |
| Scan technique | 3D MPRAGE | 3D MPRAGE |  | 3D MPRAGE |
| Number of signals (repetitions) | 1 | 2 |  | 2 |
| FOV (RL / AP / FH; mm.) | 160×256×256 | 188×240×220 |  | 188×240×220 |
| Voxel size (mm.) | 1×1×1 | 1×1×1 |  | 1×1×1 |
| TR / TE (millisec.) | 8.1 / 3.7 | 8.5 / 3.9 |  | 8.5 / 3.9 |
| Water-fat shift (pix.) | 2.268 | 2.268 |  | 2.268 |
| Bandwidth (Hz./pix.) | 191.5 | 191.5 |  | 191.5 |
| Flip angle (deg.) | 8 | 8 |  | 8 |
| Phase accell. factor (SENSE) | 1.5 (RL) | 2.5 (RL) / 2 (FH) |  | 2.5 (RL) / 2 (FH) |
| Acquisition direction | Sagittal | Axial |  | Axial |
| Duration | 5 min 58 sec | 6 min 3 sec |  | 6 min 3 |
| **Functional (BOLD) MRI** |  | *MB scans* | *Seq. scans* |  |
| Number of scans | 1 | 2 | 3 | 4 |
| Scan technique | GE-EPI | GE-EPI | GE-EPI | GE-EPI |
| FOV (RL / AP / FH) | 138×192×192 | 240×240×118 | 240x240x122 | 240x240x122 |
| Voxel size (mm.) | 3×3×3 | 3×3×3 | 3×3×3 | 3×3×3 |
| Matrix size | 64x64 | 80x80 | 80x80 | 80x80 |
| Nr. of slices | 40 | 36 | 37 | 37 |
| Slice gap (mm.) | 0.3 | 0.3 | 0.3 | 0.3 |
| TR / TE (ms.) | 2200 / 28 | 750 / 28 | 2000 | 28 |
| Water-fat shift (pix.) | 12.481 | 11.001 | 11.502 | 11.502 |
| Bandwidth (Hz/Pix) | 34.6 | 39.5 | 37.8 | 37.8 |
| Flip angle (deg.) | 90 | 60 | 76.1 | 76.1 |
| Phase accell. factor (SENSE) | 0 | 2 (AP) | 2 (AP) | 2 (AP) |
| Phase encoding direction | P >> A | P >> A | P >> A | P >> A |
| Slice encoding direction | L >> R | F >> H | F >> H | F >> H |
| Nr. of dummy scans | 2 | 2 | 2 | 2 |
| Dynamic stabilization | none | enhanced | enhanced | enhanced |
| Duration | 10 min 38 sec | variable | variable | variable |
| ***Diffusion-weighted images*** |  |  |  |  |
| Number of scans | 3 | 1 | 1 | 1 |
| Scan technique | SE-DWI | SE-DWI | SE-DWI | SE-DWI |
| Nr. of b0 images | 1 | 1 | 1 | 1 |
| Nr. of diffusion-weighted dirs. | 32 | 32 | 32 | 32 |
| Sampling scheme | Half sphere | Half sphere | Half sphere | Half sphere |
| DWI b-value | 1000 s/mm<sup>2</sup> | 1000 s/mm<sup>2</sup> | 1000 s/mm<sup>2</sup> | 1000 s/mm<sup>2</sup> |
| Voxel size (mm.) | 2×2×2 | 2×2×2 | 2×2×2 | 2×2×2 |
| FOV (RL / AP / FH) | 224×224×120 | 224×224×120 | 224×224×120 | 224×224×120 |
| Matrix size | 112×112 | 112×112 | 112×112 | 112×112 |
| Nr. of slices | 60 | 60 | 60 | 60 |
| Slice gap (mm.) | 0 | 0 | 0 | 0 |
| TR / TE | 6370 / 75 | 7456 / 86 | 7456 / 86 | 7456 / 86 |
| Water-fat shift (pix.) | 12.861 | 18.926 | 18.926 | 18.926 |
| Bandwidth (Hz/pix.) | 33.8 | 22.9 | 22.9 | 22.9 |
| Flip angle (deg.) | 90 | 90 | 90 | 90 |
| Phase accell. (SENSE) | 3 (AP) | 2 (AP) | 2 (AP) | 2 (AP) |
| Phase encoding dir. | P >> A | P >> A | P >> A | P >> A |
| Slice encoding direction | F >> H | F >> H | F >> H | F >> H |
| Duration | 4 min 49 sec | 5 min 27 sec | 5 min 27 sec | 5 min 27 sec |

During functional MRI scans, additional physiological data was recorded. Respiratory traces were recorded using a respiratory belt (air filled cushion) bound on top of the subject’s diaphragm using a velcro band. Cardiac traces were recorded using a plethysmograph attached to the subject’s left ring finger. Data was transferred to the scanner PC as plain-text files (Philips “SCANPHYSLOG” files) using a wireless recorder with a sampling frequency of 496 Hz.

Experimental paradigms for the functional MRI runs were shown on a 61 × 36 cm screen using a DLP projector with a 60 Hz refresh rate (ID1000 and PIOP1) or on a Cambridge Electronics BOLDscreen 32 IPS LCD screen with a 120 Hz refresh rate (PIOP2), both placed at 113 cm distance from the mirror mounted on top of the head coil. Sound was presented via a MRConfon sound system. Experimental tasks were programmed using Neurobs Presentation (Neurobehavioral Systems Inc, Berkeley, U.S.A.) and run on a Windows computer with a dedicated graphics card. To allow subject responses in experimental tasks (PIOP1 and PIOP2 only), participants used MRI-compatible fibre optic response pads with four buttons for each hand (Cambridge Research Systems, Rochester, United Kingdom).

### ID1000 specifics

In this section, we describe the subject recruitment, subject characteristics, data collection protocol, and functional MRI paradigm of the ID1000 study.

#### Subjects

The data from the ID1000 sample was collected between 2010 and 2012. The faculty’s ethical committee approved this study before data collection started (EC number: 2010-BC-1345). Prior to the experiment, subjects were informed about the goal and scope of the research, the MRI procedure, safety measures, general experimental procedures, privacy and data sharing concerns, and voluntary nature of the project (i.e., subjects were told that they could stop with the experiment at any time, without giving a reason for it).

Before the start of the experiment, subjects signed an informed consent form and were screened for MRI safety. We recorded data from 992 subjects of which 928 are included in the dataset (see Technical validation for details on the exclusion procedure). Subjects were recruited through a recruitment agency (Motivaction International B.V.) in an effort to get a sample that was representative of the general Dutch population in terms of educational level (as defined by the Dutch government^13^), but drawn from only a limited age range (19 to 26; see Table 4 for details). We chose this limited age range to minimize the effect of aging on any brain-related covariates. A more detailed description of educational level and other demographic variables can be found in the section “Subject variables”.

#### Data collection protocol

Before coming to the scan center, subjects completed a questionnaire on background information (to determine educational level, which was used to draw a representative sample). When invited to participate, subjects completed, at the scan center, an extensive set of questionnaires and tests, including a general demographics questionnaire, the Intelligence Structure Test^14^, the “trait” part of the State-Trait Anxiety Inventory (STAI)^15^, a behavioral avoidance/inhibition questionnaire (BIS/BAS)^16^, multiple personality questionnaires — amongst them the MPQ^17^ and the NEO-FFI^18,19^ and several behavioral tasks. The psychometric variables of the tests included in the current dataset are described in the section “Psychometric variables”.

Testing took place from 9 AM until 4 PM and on each day two subjects were tested. One subject began with the IST intelligence test, while the other subject started with the imaging part of the experiment. For the MRI part, we recorded three T1-weighted scans, three diffusion-weighted scans, and one functional (BOLD) MRI scan (in that order). Afterwards, the subjects switched and completed the other part. After these initial tasks, the subjects participated in additional experimental tasks, some of which have been reported in other publications^20,21^ and are not included in this dataset.

#### Functional MRI paradigm

During functional MRI acquisition, subjects viewed a movie clip consisting of a (continuous) compilation of 22 natural scenes taken from the movie Koyaanisqatsi^22^ with music composed by Philip Glass. The scenes varied in length from approximately 5 to 40 seconds with “cross dissolve” transitions between scenes. The movie clip extended 16 degrees visual angle (resolution 720×576, movie frame rate of 25 Hz). The scenes were selected because they sample a set of visual parameters (textures and objects with different sizes and different rates of movements) broadly. The onset of the movie clip was triggered by the first volume of the fMRI acquisition and had a duration of 11 minutes (which is slightly longer than the fMRI scan, i.e., 10 minutes and 38 seconds). The movie clip is available in the “stimuli” subdirectory of the ID1000 dataset (with the filename *task-moviewatching_desc-koyaanisqatsi_movie.mp4*).

### PIOP1 and PIOP2 specifics

In this section, we describe the subject recruitment, subject characteristics, data collection protocol, and functional MRI paradigm of the PIOP1 and PIOP2 studies. These two studies are described in a common section because their data collection protocols were very similar.

#### Subjects

Data from the PIOP1 dataset were collected between May 2015 and April 2016 and data from the PIOP2 dataset between March 2017 and July 2017. The faculty’s ethical committee approved these studies before data collection started (PIOP1 EC number: 2015-EXT-4366, PIOP2 EC number: 2017-EXT-7568). Data was recorded from 248 subjects (PIOP1) and 242 subjects (PIOP2), of which 216 (PIOP1) and 226 (PIOP2) are included in AOMIC (see Technical validation for details on the exclusion procedure). Subjects were all university students (from the Amsterdam University of Applied Sciences or the University of Amsterdam) recruited through the University websites, posters placed around the university grounds, and Facebook. A description of demographic and other subject-specific variables can be found in the section subject variables.

#### Data collection protocol

Prior to the research, subjects were informed about the goal of the study, the MRI procedure and safety, general experimental procedure, privacy and data sharing issues, and the voluntary nature of participation through an information letter. Each testing day (which took place from 8.30 AM until 1 PM), four subjects were tested. First, all subjects filled in an informed consent form and completed an MRI screening checklist. Then, two subjects started with the MRI part of the experiment, while the other two completed the demographic and psychometric questionnaires (described below) as well as several tasks that are not included in AOMIC.

The MRI session included a survey scan, followed by a T1-weighted anatomical scan. Then, several functional MRI runs (described below) and a single diffusion-weighted scan were recorded. Details about the scan parameters can be found in Table 2.

#### Functional MRI paradigms

In this section, we will describe the experimental paradigms used during fMRI acquisition. See Figure 2 for a visual representation of each paradigm.

**Figure 2.**
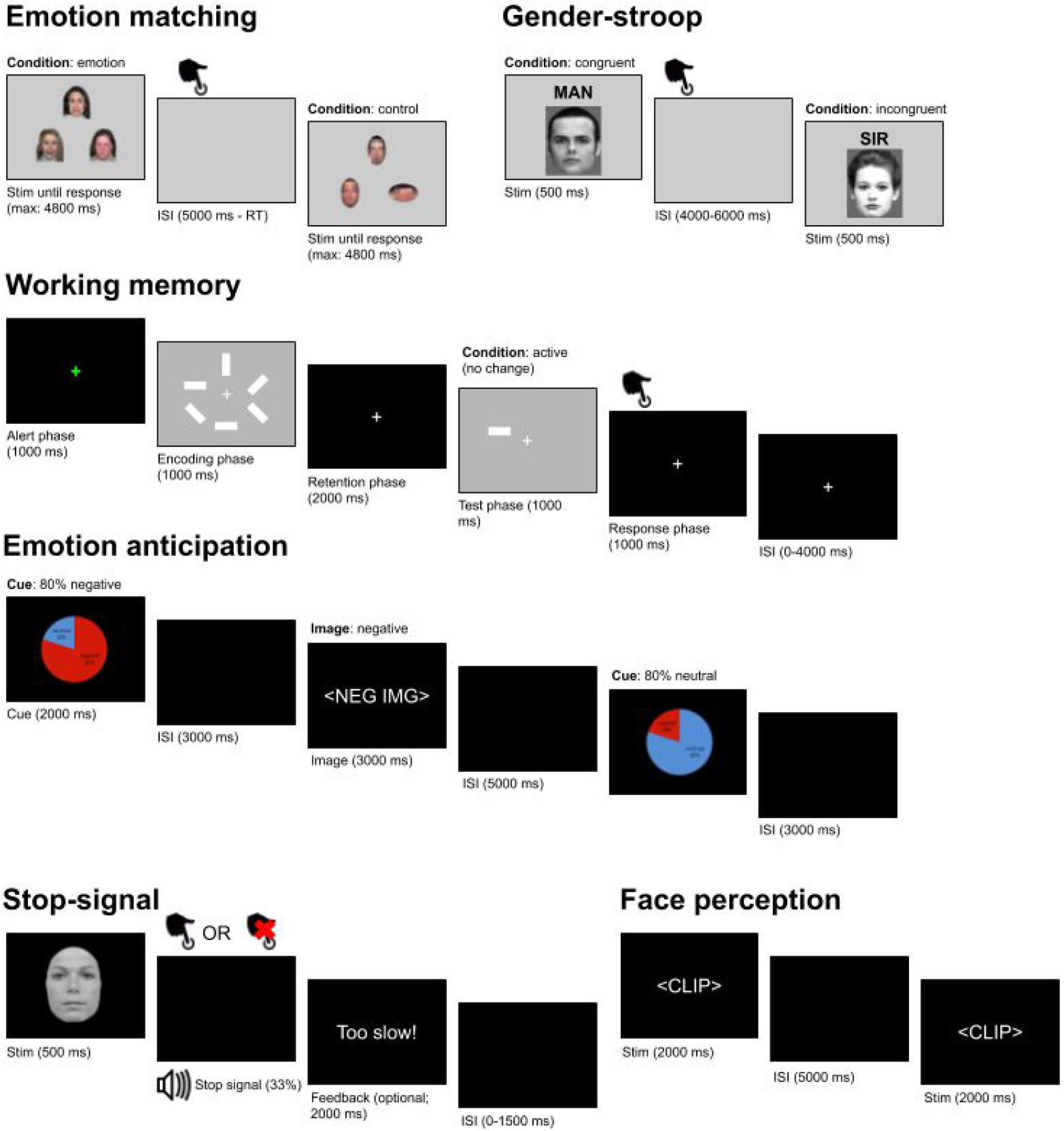
A visual representation of all experimental paradigms during task-based fMRI. ISI: inter-stimulus interval.

##### Emotion matching (PIOP1+2)

The goal of the “emotion matching” task is to measure processes related to (facial) emotion processing. The paradigm we used was based on a study by Hariri and colleagues (2000)^23^. In each trial, subjects were presented with either color images of an emotional target face (top) and two emotional probe faces (bottom left and bottom right; “emotion” condition) or a target oval (top) and two probe ovals (bottom left and bottom right; “control” condition) on top of a gray background (RGB: 248, 248, 248) and were instructed to either match the emotional expression of the target face (“emotion” condition) or the orientation or the target oval (“control” condition) as quickly as possible by pushing a button with the index finger of their left or right hand. The target and probes disappeared when the subject responded (or after 4.8 seconds). A new trial always appeared 5 seconds after the onset of each trial. In between the subject’s response and the new trial, a blank screen was shown. Trials were presented in alternating “control” and “emotion” blocks consisting of six stimuli of 5 seconds each (four blocks each, i.e., 48 stimuli in total). Stimuli always belonged to the same block, but the order of stimuli within blocks was randomized across participants.

The faces always displayed either stereotypical anger or fear. Within trials, always exactly two faces portrayed the same expression. Both male and female faces and white, black, and Asian faces were used, but within a single trial, faces were always of the same sex and ethnicity category (white or Asian/black). Face pictures were derived from the NimStim Face Stimulus set^24^. The oval stimuli were created by pixelating the face stimuli and were approximately the same area as the face stimuli (making them color and size matched to the face stimuli) and were either presented horizontally (i.e., the long side was horizontally aligned) or vertically (i.e., the long side was vertically aligned). Within trials, always exactly two ovals were aligned in the same way.

The fMRI “event files” (with the identifier *_events*) associated with this task contain information of the trial onset (the moment the faces/ovals appeared on screen, in seconds), duration (how long the faces/ovals were presented, in seconds), trial type (either “control” or “emotion”), response time (how long it took the subject to respond, logged “n/a” in case of no response), response hand (either “left”, “right”, or “n/a” in case of no response), response accuracy (either “correct”, “incorrect”, or “miss”), orientation to match (either “horizontal”, “vertical”, or “n/a” in case of emotion trials), emotion match (either “fear”, “anger”, or “n/a” in case of control trials), gender of the faces (either “male”, “female”, of “n/a” in case of control trials), and ethnicity of the target and probe faces (either “caucasian”, “asian”, “black”, or “n/a” in case of control trials).

##### Working memory task (PIOP1+2)

The goal of the working memory task was to measure processes related to visual working memory. The paradigm we used was based on Pessoa and colleagues (2002)^25^. Trials belong to one of three conditions: “active (change)”, “active (no change)”, or “passive”. Each trial consists of six phases: an alert phase (1 second), an encoding phase (1 second), a retention phase (2 seconds), a test phase (1 second), a response phase (1 second), and an inter-stimulus interval (0-4 seconds). Subjects were instructed to keep focusing on the fixation sign, which was shown throughout the entire trial, and completed a set of practice trials before the start of the actual task.

In all trial types, trials started with an alert phase: a change of color of the fixation sign (a white plus sign changing to green, RGB [0, 255, 0]), lasting for 1 second. In the encoding phase, for “active” trials, an array of six white bars with a size of 2 degrees visual angle with a random orientation (either 0, 45, 90, or 135 degrees) arranged in a circle was presented for 1 second. This phase of the trial coincided with a change in background luminance from black (RGB: [0, 0, 0]) to gray (RGB: [120, 120, 120]). For “passive” trials, only the background luminance changed (but no bars appeared) in the encoding phase. In the subsequent retention phase, for all trial types, a fixation cross was shown and the background changed back to black, lasting 2 seconds. In the test phase, one single randomly chosen bar appeared (at one of the six locations from the encoding phase) which either matched the original orientation (for “active (no change)” trials) or did not match the original orientation (for “active (change)” trials), lasting for 1 second on a gray background. For “passive” trials, the background luminance changed and, instead of a bar, the cue “respond left” or “respond right” was shown in the test phase. In the response phase, lasting 1 second, the background changed back to black and, for “active” trials, subjects had to respond whether the array changed (button press with right index finger) or did not change (button press with left index finger). For “passive” trials, subjects had to respond with the hand that was cued in the test phase. In the inter-stimulus interval (which varied from 0 to 4 seconds), only a black background with a fixation sign was shown.

In total, there were 8 “passive” trials, 16 “active (change)” and “active (no change)” trials, in addition to 20 “null” trials of 6 seconds (which are equivalent to an additional inter-stimulus interval of 6 seconds). The sequence trials was, in terms of conditions (active, passive, null) exactly the same for all participations in both PIOP1 and PIOP2 and was optimized for BOLD-response shape estimation efficiency, but which bar or cue was shown in the test phase was chosen randomly.

The fMRI event files associated with this task contain information of the trial onset (the moment when the alert phase started, in seconds), duration (from the alert phase up to and including the response phase, i.e., always 6 seconds), trial type (either “active (change)”, “active (no change)”, or “passive”; “null” trials were not logged), response time (how long it took the subject to respond, “n/a” in case of no response), response hand (either “left”, “right”, or “n/a” in case of no response), and response accuracy (either “correct”, “incorrect”, or “miss”). Note that, in order to model the response to one or more phases of the trial, the onsets and durations should be adjusted accordingly (e.g., to model the response to the retention phase, add 2 seconds to all onsets and change the duration to 2 seconds).

##### Resting state (PIOP1+2)

During the resting state scans, participants were instructed to keep their gaze fixated on a fixation cross in the middle of the screen with a gray background (RGB: [150, 150, 150]) and to let their thoughts run freely. Eyetracking data was recorded during this scan but is not included in this dataset. The resting state scans lasted 6 minutes (PIOP1) and 8 minutes (PIOP2).

##### Face perception (PIOP1)

The face perception task was included to measure processes related to (emotional) facial expression perception. In each trial, subjects passively viewed dynamic facial expressions (i.e., short video clips) taken from the Amsterdam Facial Expression Set (ADFES)^26^, which displayed either anger, contempt, joy, or pride, or no expression (“neutral”). Each clip depicted a facial movement from rest to a full expression corresponding to one of the four emotions, except for “neutral” faces, which depicted no facial movement. All clips lasted 2 seconds and contained either North-European or Mediterranean models, all of whom were female. After each video, a fixed inter-stimulus interval of 5 seconds followed. Each emotional facial expression (including “neutral”) was shown 6 times (with different people showing the expression each time), except for one, which was shown 9 times. Which emotional expression (or “neutral”) was shown an extra three times was determined randomly for each subject. Importantly, the three extra presentations always contained the same actor and were always presented as the first three trials. This was done in order to make it possible to evaluate the possible effects of stimulus repetition.

The fMRI event files associated with this task contain information of the trial onset (the moment when the clip appeared on screen, in seconds), duration (of the clip, i.e., always 2 seconds), trial type (either “anger”, “joy”, “contempt”, “pride”, or “neutral”), sex of the model (all “female”), ethnicity of the model (either “North-European” or “Mediterranean”), and the ADFES ID of the model.

##### Gender-stroop task (PIOP1 only)

The goal of the gender-stroop task was to measure processes related to cognitive conflict and control^27^ (see also ref^28^ for an investigation of these processes using the PIOP1 gender-stroop data). We used the face-gender variant of the Stroop task (which was adapted from ref^29^), often referred to as the “gender-stroop” task. In this task, pictures of twelve different male and twelve different female faces are paired with the corresponding (i.e., congruent) or opposite (i.e., incongruent) label. For the labels, we used the Dutch words for “man”, “sir”, “woman”, and “lady” using either lower or upper case letters. The labels were located just above the head of the face.

On each trial, subjects were shown a face-label composite on top of a gray background (RGB: [105, 105, 105]) for 0.5 seconds, which was either “congruent” (same face and label gender) or “incongruent” (different face and label gender). Stimulus presentation was followed by an inter-stimulus interval ranging between 4 and 6 seconds (in steps of 0.5 seconds). Subjects were always instructed to respond to the gender of the pictured face, ignoring the distractor word, as fast as possible using their left index finger (for male faces) or right index finger (for female faces). There were 48 stimuli for each condition (“congruent” and “incongruent”).

The fMRI event files associated with this task contain information of the trial onset (the moment when the face-label composite appeared on screen, in seconds), duration (of the face-label composite, i.e., always 0.5 seconds), trial type (either “incongruent” or “congruent”), gender of the face (either “male or “female”), gender of the word (either “male” or “female”), response time (in seconds), response hand (either “left”, “right”, or “n/a” in case of no response), response accuracy (either “correct”, “incorrect”, or “miss”).

We note that, due to an absence of any “baseline” (or “null”) trials, the condition-regressors (related to congruent and incongruent trials) are strongly negatively correlated (between −0.8 and −0.95), which lead to inefficient first-level GLM estimation and consequently underpowered (i.e., high-variance) parameter estimates. Other factors, such as response accuracy, however, are much less correlated and are efficiently estimated (see, e.g., Figure 6 for the result of a group-level whole-brain analysis of the “incorrect > correct” contrast).

##### Emotion anticipation task (PIOP1 only)

We included the emotion anticipation task to measure processes related to (emotional) anticipation and curiosity. The paradigm was based on paradigms previously used to investigate (morbid) curiosity^30,31^. In this task, subjects viewed a series of trials containing a cue and an image. The cue could either signal an 80% chance of being followed by a negatively valenced image (and a 20% chance of a neutral image) or an 80% chance of being followed by a neutral image (and a 20% change of a negatively valenced image). The cue was shown on top of a black background for 2 seconds, followed by a fixed interval of 3 seconds. After this interval, either a negative or neutral image was shown for 3 seconds, with a frequency that corresponds to the previously shown cue. In other words, for all trials with, for example, a cue signalling an 80% chance of being followed by a neutral image, it was in fact followed by a neutral image in 80% of the times. After the image, a fixed inter-stimulus interval of five seconds followed. In total, 15 unique negative images and 15 unique neutral images were shown, of which 80% (i.e., 12 trials) was preceded by a “valid” cue. Which stimuli were paired with valid or invalid cues was determined randomly for each subject. The order of the trials, given the four possible combinations (valid cue + negative image, invalid cue + negative image, valid cue + neutral image, invalid cue + neutral image), was drawn randomly from one of four possible sequences, which were generated using OptSeq (https://surfer.nmr.mgh.harvard.edu/optseq/) to optimize the chance of finding a significant interaction between cue type and image valence.

The cue was implemented as a pie chart with the probability of a neutral image in blue and the probability of the negative image in red with the corresponding labels and probabilities (e.g., “negative 20%”) superimposed for clarity. The images were selected from the IAPS database^32^ and contained images of mutilation, violence, and death (negative condition) and of people in neutral situations (neutral condition).

The fMRI event files associated with this task contain information on the trial onset (the moment when either the cue or image appeared on the screen, in seconds), duration (of either the cue or image), and trial type. Trial type was logged separately for the cues (“negative”, indicating 80% probability of a negative image, and “neutral”, indicating 80% probability of a neutral image) and images (“negative” or “neutral”).

##### Stop-signal task (PIOP2 only)

The stop-signal task was included to measure processes related to response inhibition. This specific implementation of the stop-signal paradigm was based on Jahfari et al. (2015)^33^. Subjects were presented with trials (*N* = 100) in which an image of either a female or male face (chosen from 9 exemplars) was shown for 500 ms on a black background. Subjects had to respond whether the face was female (right index finger) or male (left index finger) as quickly and accurately as possible, except when an auditory stop signal (a tone at 450 Hz for 0.5 seconds) was presented (on average 33% of the trials). The delay in presentation of the stop signal (i.e., the “stop signal delay”) was at start of the experiment 250 milliseconds, but was shortened with 50 ms if stop performance, up to that point, was better than 50% accuracy and shortened with 50 ms if it was worse. Each trial had a duration of 4000 ms and was preceded by a jitter interval (0, 500, 1000 or 1500 ms). If subjects responded too slow, or failed to respond an additional feedback trial of 2000 ms was presented. Note that due to this additional feedback trial and the fact that subjects differed in how many feedback trials they received, the fMRI runs associated with this task differ in length across subjects (i.e., the scan was manually stopped after 100 trials). Additionally 10% (on average) null trials with a duration of 4000 ms were presented randomly.

The fMRI event files associated with this task contain information on the trial onset (the moment when a face was presented, in seconds), duration (always 0.5083 seconds), trial type (go, succesful_stop, unsuccesful_stop), if and when a stop signal was given (in seconds after stimulus onset), if and when a response time was given (in seconds after stimulus onset), response of the subject (left, right), and the sex of the image (male, female).

### Subject variables (all datasets)

In AOMIC, several demographic and other subject-specific variables are included per dataset. Below, we describe all variables in turn (note that some variables are not included in all datasets, see Table 5).

#### Age

We asked subjects for their date of birth at the time they participated. From this, we computed their age rounded to the nearest quartile (for privacy reasons). See Table 3 for descriptive statistics of this variable. See Table 3 for descriptive statistics of this variable.

**Table 3.**
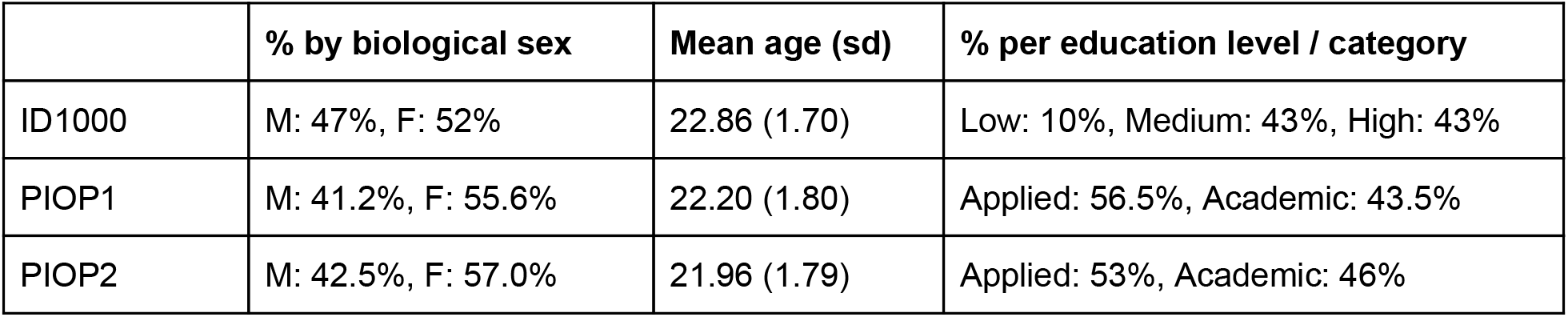
Descriptive statistics for biological sex, age, and education level for all three datasets.

|  | % by biological sex | Mean age (sd) | % per education level / category |
| --- | --- | --- | --- |
| ID1000 | M: 47%, F: 52% | 22.86 (1.70) | Low: 10%, Medium: 43%, High: 43% |
| PIOP1 | M: 41.2%, F: 55.6% | 22.20 (1.80) | Applied: 56.5%, Academic: 43.5% |
| PIOP2 | M: 42.5%, F: 57.0% | 21.96 (1.79) | Applied: 53%, Academic: 46% |

#### Biological sex and gender identity

In all three studies, we asked subjects for their biological sex (of which the options were either male or female). For the ID1000 dataset, after the first 400 subjects, we additionally asked to what degree subjects felt male and to what degree they felt female (i.e., gender identity; separate questions, 7 point likert scale, 1 = not at all, 7 = strong). The exact question in Dutch was: ‘ik voel mij een man’, ‘ik voel mij een vrouw’. This resulted in 0.3% of subjects scoring opposite on gender identity compared to their biological sex and 92% of females and 90% of males scoring conformable with their sex.

#### Sexual orientation

For the ID1000 dataset, after the first 400 subjects, we additionally asked to what degree subjects were attracted to men and women (both on a 7 point likert scale, 1 = not at all, 7 = strong). The exact question in Dutch was: ‘ik val op mannen’ and ‘ik val op vrouwen’.

Of the 278 subjects with a male sex 7,6% indicated to be attracted to men (score of 4 or higher), of the 276 subjects with a female sex 7,2% indicated to be attracted to women (score of 4 or higher). Of the 554 subjects who completed these questions 0.4% (n=2) indicated not be attracted to either men or women and 0.9% indicated to be strongly attracted to both men and women.

#### BMI

Subjects were asked for (PIOP1 and PIOP2) or we measured (ID1000) subjects’ height and weight on the day of testing, from which we calculated their body-mass-index (BMI), which we rounded to the nearest integer. Note that height and weight are not included in AOMIC (for privacy reasons), but BMI is.

#### Handedness

Subjects were asked for their dominant hand (options were “left”, “right”). For PIOP1 and PIOP2 we also included the option “both”.

#### Educational level / category

Information about subjects’ educational background is recorded differently for ID1000 and the PIOP datasets, so they are discussed separately. Importantly, while we included data on educational *level* for the ID1000 dataset, we only include data on educational *category* for the PIOP datasets because they contain little variance in terms of educational level.

##### Educational level (ID1000)

As mentioned, for the ID1000 dataset we selected subjects based on their educational level in order to achieve a representative sample of the Dutch population (on that variable). We did this by asking for their highest *completed* educational level. In AOMIC, however, we report the educational level (three point scale: low, medium, high) on the basis of the completed *or* current level of education (which included the scenario in which the subject was still a student), which we believe reflects educational level for our relatively young (19-26 year old) better. Note that this difference in criterion causes a substantial skew towards a higher educational level in our sample relative to the distribution of educational level in the Dutch population (see Table 4).

**Table 4.** Distribution of educational level in the Dutch population (in 2010) and in our ID1000 sample. The data from the education level of subjects’ parents was used to compute background socio-economic status.

| **Educational level** | *Population* | *Subjects* | *Father* | *Mother* |
| --- | --- | --- | --- | --- |
| Low | 35% | 10% | 36% | 32% |
| Medium | 39% | 43% | 27% | 24% |
| High | 26% | 47% | 37% | 44% |

##### Educational category (PIOP)

Relative to ID1000, there is much less variance in educational level within the PIOP datasets as these datasets only contain data from university students. As such, we only report whether subjects were, at the time of testing, studying at the Amsterdam University of Applied Sciences (category: “applied”) or at the University of Amsterdam (category: “academic”).

##### Background socio-economic status (SES)

In addition to reporting their own educational level, subjects also reported the educational level (see Table 4) of their parents and the family income in their primary household. Based on this information, we determined subjects’ background social economical status (SES) by adding the household income — defined on a three point scale (below modal income, 25%: 1, between modal and 2 × modal income, 57%: 2, above 2 × modal income, 18%: 3) — with the average educational level of the parents — defined on a three point scale (low: 1, medium: 2, high: 3). This revealed that, while the educational level of the subjects is somewhat skewed towards “high”, SES is well distributed across the entire spectrum (see Table 5).

**Table 5.** Distribution of background SES.

| Background SES | % of subjects |
| --- | --- |
| 2-3 | 16% |
| 3-4 | 26% |
| 4-5 | 28% |
| 5-6 | 19% |
| >6 | 11% |

#### Religion (PIOP1 and ID1000 only)

For both the PIOP1 and ID1000 datasets, we asked subjects whether they considered themselves religious, which we include as a variable for these datasets (recoded into the levels “yes” and “no”). Of the subjects that participated in the PIOP1 study 18.0% indicated to be religious, for the subjects in the ID1000 projects this was 21.2%. For the ID1000 dataset we also asked subjects if they were raised religiously (N = 928, 34.1%) and to what degree religion played a daily role in their lives (in Dutch, “Ik ben dagelijks met mijn geloof bezig”, 5 point likert scale, 1 = not at all applicable, 5 = very applicable).

### Psychometric variables (all datasets)

#### BIS/BAS (ID1000 only)

The BIS/BAS scales are based on the idea that there are two general motivational systems underlying behavior and affect: a behavioral activation system (BAS) and a behavioral inhibition system (BIS). The scales of the BIS/BAS attempt to measure these systems^34^. The BAS is believed to measure a system that generates positive feedback while the BIS is activated by conditioned stimuli associated with punishment.

The BIS/BAS questionnaire consists of 20 items (4 point scale). The BIS scale consists of 7 items. The BAS scale consists of 13 items and contains three subscales, related to impulsivity (BAS-Fun, 4 items), reward responsiveness (BAS-Reward, 5 items) and the pursuit of rewarding goals (BAS-Drive, 4 items).

#### STAI-T (ID1000)

We used the STAI^35,36^ to measure trait anxiety (STAI-*T*). The questionnaire consists of two scales (20 questions each) that aim to measure the degree to which anxiety and fear are a trait and part of the current state of the subject; subjects only completed the trait part of the questionnaire, which we include in the ID1000 dataset.

#### NEO-FFI (all datasets)

The NEO-FFI^18,37^ is a Big 5 personality questionnaire that consists of 60 items (12 per scale). It measures neuroticism (“NEO-N”), extraversion (“NEO-E”), openness to experience (“NEO-O”), agreeableness (“NEO-A”), and conscientiousness (“NEO-C”). Neuroticism is the opposite of emotional stability, central to this construct is nervousness and negative emotionality. Extraversion is the opposite of introversion, central to this construct is sociability — the enjoyment of others’ company. Openness to experience is defined by having original, broad interests, and being open to ideas and values. Agreeableness is the opposite of antagonism. Central to this construct are trust, cooperation and dominance. Conscientiousness is the opposite of un-directedness. Adjectives associated with this construct are thorough, hard-working and energetic.

#### IST (ID1000 only)

The Intelligence Structure Test (IST)^14^ is an intelligence test measuring crystallized intelligence, fluid intelligence, and memory, through tests using verbal, numerical, and figural information. The test consists of 590 items. The three measures (crystallized intelligence, fluid intelligence, and memory) are strongly positively correlated (between *r* = .58 and *r* = .68, see Table 9) and the sum score of these values form the variable “total intelligence”.

#### Raven’s matrices (PIOP only)

As a proxy for intelligence, subjects performed the 36 item version (set II) of the Raven’s Advanced Progressive Matrices Test^38,39^. We included the sum-score (with a maximum score of 36) in the PIOP datasets.

Importantly, all subject variables and psychometric variables are stored in the *participants.tsv* file in each study’s data repository. In Table 6, all variables and associated descriptions are listed for convenience.

**Table 6.** Description of the subject variables and psychometric variables contained in AOMIC. The “variable name” coincides with the column name for that variable in the *participants.tsv* file. Note that missing values in this file are coded with “n/a”. A.U.: arbitrary units.

| Variable name | Description | Levels / range | Units | Included in |
| --- | --- | --- | --- | --- |
| age | Subject age at day of measurement | - | Quantiles | All |
| sex | (Biological) sex | male, female | - | All |
| gender_identity_F | To what degree the subject felt female | 1-7 | A.U. | ID1000 |
| gender_identity_M | To what degree the subject felt male | 1-7 | A.U. | ID1000 |
| sexual_attraction_M | To what degree the subject was attracted to males | 1-7 | A.U. | ID1000 |
| sexual_attraction_F | To what degree the subject was attracted to females | 1-7 | A.U. | ID1000 |
| BMI | Body-mass-index | - | kg/m<sup>2</sup> | All |
| handedness | Dominant hand | left, right |  | ID1000 |
| handedness | Dominant hand | left, right, ambidextrous | - | PIOP1&2 |
| background_SES | Social economical status of family in which the subject was raised. | 2-6 | A.U. | ID1000 |
| educational_level | Highest achieved (or current) educational level | low, medium, high | - | ID1000 |
| educational_category | Highest achieved (or current) education type | applied, academic | - | PIOP1&2 |
| religious | Whether the subject is religious or not | no, yes | - | PIOP1 |
| religious_upbringing | Whether the subject was raised religiously or not | no, yes | - | ID1000 |
| religious_now | Whether the subject is religious now | no, yes | - | ID1000 |
| religious_importance | To what degree religion plays a role in the subject's daily life | 1-5 | A.U. | ID1000 |
| IST_fluid | IST fluid intelligence subscale | - | A.U. | ID1000 |
| IST_memory | IST memory subscale | - | A.U. | ID1000 |
| IST_crystallised | IST crystallised intelligence subscale | - | A.U. | ID1000 |
| IST_total_intelligence | IST total intelligence scale | - | A.U. | ID1000 |
| BAS_drive | BAS drive scale | - | A.U. | ID1000 |
| BAS_fun | BAS fun scale | - | A.U. | ID1000 |
| BAS_reward | BAS reward scale | - | A.U. | ID1000 |
| BIS | BIS scale | - | A.U. | ID1000 |
| NEO_N | Neuroticism scale (sum score) | - | A.U. | All |
| NEO_E | Extraversion scale (sum score) | - | A.U. | All |
| NEO_O | Openness scale (sum score) | - | A.U. | All |
| NEO_A | Agreeableness scale (sum score) | - | A.U. | All |
| NEO_C | Conscientiousness scale (sum score) | - | A.U. | All |
| STAI_T | Trait anxiety (from the STAI) (sum score) | - | A.U. | ID1000 |

### Data standardization, preprocessing, and derivatives

In this section, we describe the data curation and standardization process as well as the preprocessing applied to the standardized data and the resulting “derivatives” (see Figure 3 for a schematic overview). This section does not describe this process separately for each dataset, because they are largely identically standardized and (pre)processed. Exceptions to this will be explicitly mentioned. In this standardization process, we adhered to the guidelines outlined in the Brain Imaging Data Structure (BIDS, v1.2.2)^12^, both for the “raw” data as well as the derivatives (whenever BIDS guidelines exist for that data or derivative modality).

**Figure 3.**
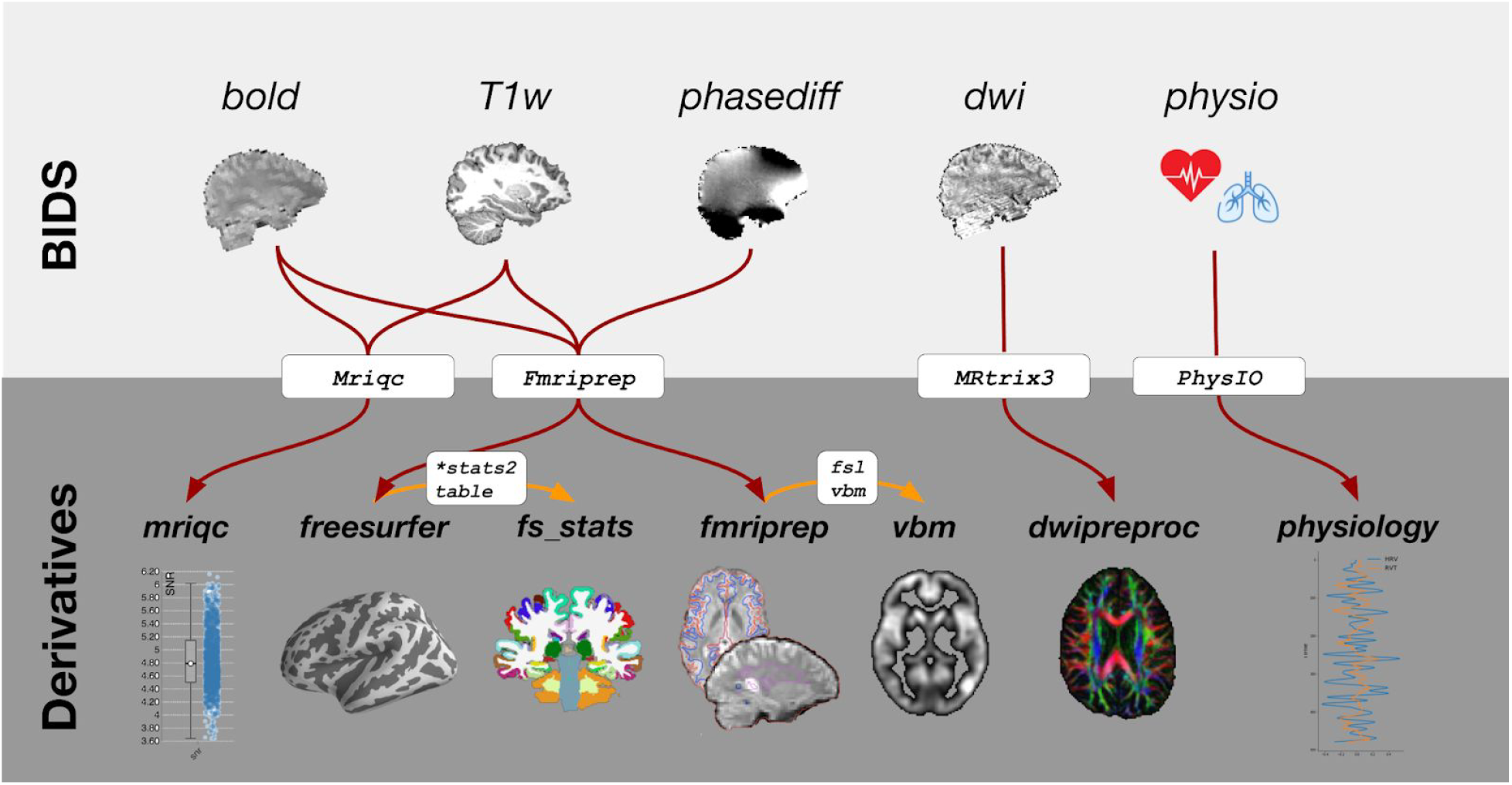
Overview of the types of data and “derivatives” included in AOMIC and the software packages used to preprocess and analyze them.

#### Raw data standardization

Before subjecting the data to any preprocessing pipeline, we converted the data to BIDS using the in-house developed package *bidsify* (see <u>Code availability</u> section for more details about the software used in the standardization process). The “BIDSification” process includes renaming of files according to BIDS convention, conversion from Philips PAR/REC format to compressed nifti, removal of facial characteristics from anatomical scans (“defacing”), and extraction of relevant metadata into JSON files.

#### Anatomical and functional MRI preprocessing

Results included in this manuscript come from preprocessing performed using Fmriprep version 1.4.1 (RRID:SCR_016216)^40,41^, a Nipype based tool (RRID:SCR_002502)^42,43^. Each T1w (T1-weighted) volume was corrected for INU (intensity non-uniformity) using N4BiasFieldCorrection v2.1.0^44^ and skull-stripped using antsBrainExtraction.sh v2.1.0 (using the OASIS template). Brain surfaces were reconstructed using recon-all from FreeSurfer v6.0.1 (RRID:SCR_001847)^45^, and the brain mask estimated previously was refined with a custom variation of the method to reconcile ANTs-derived and FreeSurfer-derived segmentations of the cortical gray-matter of Mindboggle (RRID:SCR_002438)^46^. Spatial normalization to the ICBM 152 Nonlinear Asymmetrical template version 2009c (RRID:SCR_008796)^47^ was performed through nonlinear registration with the antsRegistration tool of ANTs v2.1.0 (RRID:SCR_004757)^48^, using brain-extracted versions of both T1w volume and template. Brain tissue segmentation of cerebrospinal fluid (CSF), white-matter (WM) and gray-matter (GM) was performed on the brain-extracted T1w using fast (FSL v5.0.9, RRID:SCR_002823)^49^.

Functional data was motion corrected using mcflirt (FSL v5.0.9)^50^. "Fieldmap-less" distortion correction was performed by co-registering the functional image to the same-subject T1w image with intensity inverted^51,52^ constrained with an average fieldmap template^53^, implemented with antsRegistration (ANTs). Note that this fieldmap-less method was used even for PIOP2, which contained a phase-difference (B0) fieldmap, because we observed that Fmriprep’s fieldmap-based method led to notably less accurate unwarping than its fieldmap-less method.

Distortion-correction was followed by co-registration to the corresponding T1w using boundary-based registration^54^ with 6 degrees of freedom, using bbregister (FreeSurfer v6.0.1). Motion correcting transformations, field distortion correcting warp, BOLD-to-T1w transformation and T1w-to-template (MNI) warp were concatenated and applied in a single step using antsApplyTransforms (ANTs v2.1.0) using Lanczos interpolation.

Physiological noise regressors were extracted by applying CompCor^55^. Principal components were estimated for the two CompCor variants: temporal (tCompCor) and anatomical (aCompCor). A mask to exclude signal with cortical origin was obtained by eroding the brain mask, ensuring it only contained subcortical structures. Six tCompCor components were then calculated including only the top 5% variable voxels within that subcortical mask. For aCompCor, six components were calculated within the intersection of the subcortical mask and the union of CSF and WM masks calculated in T1w space, after their projection to the native space of each functional run. Framewise displacement^56^ was calculated for each functional run using the implementation of Nipype.

Many internal operations of Fmriprep use Nilearn (RRID:SCR_001362)^57^, principally within the BOLD-processing workflow. For more details of the pipeline see https://fmriprep.readthedocs.io/en/latest/workflows.html.

#### Diffusion MRI (pre)processing

DWI scans were preprocessed using a custom pipeline combining tools from MRtrix3 and FSL. Because we acquired multiple DWI scans per participant in the ID1000 study (but not in PIOP1 and PIOP2), we concatenated these files as well as the diffusion gradient table (*bvecs*) and b-value information (*bvals*) prior to preprocessing. Using MRtrix3, we denoised the diffusion-weighted data using *dwidenoise*^58,59^, removed Gibbs ringing artifacts using *mrdegibbs*^60^, and performed eddy current and motion correction using *dwipreproc*. Notably, *dwipreproc* is a wrapper around the GPU-accelerated (CUDA v9.1) FSL tool *eddy*^61^. Within *eddy*, we used a quadratic first-level (*--flm=quadratic*) and linear second-level model (*--slm=linear*) and outlier replacement^62^ with default parameters (*--repol*). Then, we performed bias correction using *dwibiascorrect* (which is based on *ANTs*; v2.3.1)^44^, extracted a brain mask using *dwi2mask*^63^, and corrected possible issues with the diffusion gradient table using *dwigradcheck*^64^.

After preprocessing, using MRtrix3 tools, we fit a diffusion tensor model on the preprocessed diffusion-weighted data using weighted linear least squares (with 2 iterations) as implemented in *dwi2tensor*^65^. From the estimated tensor image, a fractional anisotropy (FA) image was computed and a map with the first eigenvectors was extracted using *tensor2metric*. Finally, a population FA template was computed using *population_template* (using an affine and an additional non-linear registration).

The following files are included in the DWI derivatives: a binary brain mask, the preprocessed DWI data as well as preprocessed gradient table (*bvec*) and b-value (*bval*) files, outputs from the eddy correction procedure (for quality control purposes, see Technical validation section), the estimated parameters from the diffusion tensor model, the eigenvectors from the diffusion tensor model, and a fractional anisotropy scalar map computed from the eigenvectors. All files are named according to BIDS Extension Proposal 16 (BEP016: diffusion weighted imaging derivatives).

#### Freesurfer morphological statistics

In addition to the complete Freesurfer directories containing the full surface reconstruction per participant, we provide a set of tab-separated values (TSV) files per participant with several morphological statistics per brain region for four different anatomical parcellations/segmentations. For cortical brain regions, we used two atlases shipped with Freesurfer: the Desikan-Killiany (*aparc* in Freesurfer terms)^66^ and Destrieux (*aparc.a2009* in Freesurfer terms)^67^ atlases. For these parcellations, the included morphological statistics are volume in mm^3^, area in mm^2^, thickness in mm, and integrated rectified mean curvature in mm^−1^. For subcortical and white matter brain regions, we used the results from the subcortical segmentation (*aseg* in Freesurfer terms) and white matter segmentation (*wmparc* in Freesurfer terms) done by Freesurfer. For these parcellations, the included morphological statistics are volume in mm^3^ and average signal intensity (arbitrary units). The statistics were extracted from the Freesurfer output directories using the Freesufer functions *asegstats2table* and *aparcstats2table* and further formatted using custom Python code. The TSV files (and accompanying JSON metadata files) are formatted according to BIDS Extension Proposal 11 (BEP011: structural preprocessing derivatives).

#### Voxel-based morphology (VBM)

In addition to the Fmriprep-preprocessed anatomical T1-weighted scans, we also provide voxelwise gray matter volume maps estimated using voxel-based morphometry (VBM). We used a modified version of the FSL VBM pipeline (http://fsl.fmrib.ox.ac.uk/fsl/fslwiki/FSLVBM)^68^, an optimised VBM protocol^69^ carried out with FSL tools^70^. We skipped the initial brain-extraction stage (*fslvbm_1_bet*) and segmentation stage (first part of *fslvbm_3_proc*) and instead used the probabilistic gray matter segmentation file (in native space) from Fmriprep (i.e., *\*label-GM_probseg.nii.gz* files) directly. These files were registered to the MNI152 standard space using non-linear registration^71^. The resulting images were averaged and flipped along the x-axis to create a left-right symmetric, study-specific grey matter template. Second, all native grey matter images were non-linearly registered to this study-specific template and "modulated" to correct for local expansion (or contraction) due to the non-linear component of the spatial transformation.

#### Physiological noise processing

Physiology files were converted to BIDS-compatible compressed TSV files using the *scanphyslog2bids* package (see <u>Code availability</u>). Each TSV file contains three columns: the first contains the cardiac trace, the second contains the respiratory trace, and the third contains the volume onset triggers (binary, where 1 represents a volume onset). Each TSV file is accompanied by a JSON metadata file with the same name, which contains information about the start time of the physiology recording relative to the onset of the first volume. Because the physiology recording always starts before the fMRI scan starts, the start time is always negative (e.g., a start time of −42.01 means that the physiology recording started 42.01 seconds before the onset of the first volume). After conversion to BIDS, the estimated volume triggers and physiology traces were plotted, visually inspected for quality, and excluded if either of the physiology traces had missing data for more than ten seconds or if the volume triggers could not be estimated.

The physiology data was subsequently used to estimate fMRI-appropriate nuisance regressors using the *TAPAS PhysIO* package (see Code availability)^72^. Using this package, we specifically estimated 18 “RETROICOR” regressors^73^ based on a Fourier expansion of cardiac (order: 2) and respiratory (order: 3) phase and their first-order multiplicative terms (as defined in ref^74^). In addition, we estimated a heart-rate variability (HRV) regressor by convolving the cardiac trace with a cardiac response function^75^ and a respiratory volume by time (RVT) regressor by convolving the respiratory trace with a respiration response function^76^.

#### Code availability

All code used for curating, annotating, and (pre)processing AOMIC are version-controlled using git and can be found in project-specific Github repositories within the NILAB-UvA Github organization: https://github.com/orgs/NILAB-UvA. Many pre and postprocessing steps were identical across datasets, so the code for these procedures is stored in a single repository: https://github.com/NILAB-UvA/AOMIC-common-scripts. Possible parameters are all hard-coded within the scripts, except for a single positional parameter pointing to the directory to be processed. All code was developed on a Linux system running Ubuntu 16.04. For custom Python-based scripts, we used Python version 3.7.

For curation, preprocessing, and analysis of the datasets, we used a combination of existing packages and custom scripts (written in Python or bash). To convert the data to the Brain Imaging Data Structure (BIDS)^12^, we used the in-house developed, publicly available software package *bidsify* (v0.3; https://github.com/NILAB-UvA/bidsify), which in turn uses the *dcm2niix* (v1.0.20181125)^77^ to convert the Philips PAR/REC files to compressed nifti files. In contrast to the data from PIOP1 and PIOP2 (which were converted to nifti using *dcm2niix*), *r2aGUI* (v2.7.0; http://r2agui.sourceforge.net/) was used to convert the data from ID1000. Because *r2aGUI* does not correct the gradient table of DWI scans for slice angulation, we used the *angulation_correction_Achieva* Matlab script (version December 29, 2007) from Jonathan Farrell to do so (available for posterity at https://github.com/NILAB-UvA/ID1000/blob/master/code/bidsify/DTI_gradient_table_ID1000.m). To remove facial characteristics from anatomical scans, we used the *pydeface* package (v.1.1.0)^78^. Finally, to convert the raw physiology files (i.e., Philips “SCANPHYSLOG” files) to BIDS, we used the in-house developed, publicly available Python package *scanphyslog2bids* (v0.1; https://github.com/lukassnoek/scanphyslog2bids).

Anatomical and functional MRI preprocessing were done using Fmriprep (v1.4.1; see the Derivatives section for extensive information about Fmriprep’s preprocessing pipeline)^40^. For our DWI preprocessing pipeline, we used tools from the MRtrix3 package (www.mrtrix.org; v3.0_RC3)^79^ and FSL (v6.0.1)^61,70^. For the VBM and dual regression pipelines, we used FSL (v6.0.1)^80^. To create the files with Freesurfer-based metrics across all participants, we used Freesurfer version 6.0.0^81^. Physiological nuisance regressors (RETROICOR and HRV/RVT regressors) were estimated using the TAPAS PhysIO Matlab package (v3.2.0)^72^.

First-level functional MRI analyses for technical validation were implemented using the Python package *nistats* (v0.0.1b2)^57^ and *nilearn* (v0.6.2)^82^. For the inter-subject correlation analysis the Brain Imaging Analysis Kit was used (BrainIAK, http://brainiak.org, v0.10; RRID:SCR_014824)^83^. Plotting brain images was done using FSLeyes (v0.32)^84^ and plotting statistical plots was done using the Python packages *seaborn*^85^ and Matplotlib^86^.

### Data Records

#### Data formats and types

In AOMIC, the majority of the data is stored in one of four formats. First, all volumetric (i.e., 3D or 4D) MRI data is stored in compressed “NIfTI” files (NIfTI-1 version; extension: .*nii.gz*). NIfTI files contain both the data and metadata (stored in the *header*) and can be loaded into all major neuroimaging analysis packages and programming languages (e.g., using the *nibabel* package^#1^ for Python, using the *oro.nifti* package in R^#2^, and natively in Matlab version R2017b and higher). Second, surface (i.e., vertex-wise) MRI data is stored in “Gifti” files (https://www.nitrc.org/projects/gifti/; extension: .*gii*). Like NIfTI files, Gifti files contain both data and metadata and can be loaded in several major neuroimaging software packages (including Freesurfer, FSL, AFNI, SPM, and Brain Voyager) and programming languages (e.g., using the *nibabel* package for Python and the *gifti* package^#3^ for R).

Third, data organized as tables (i.e., observations in rows and properties in columns), such as physiological data and task-fMRI event log files, are stored in tab-separated values (TSV) files, which contain column names as the first row. TSV files can be opened using spreadsheet software (such as Microsoft Excel or Libreoffice Calc) and read using most major programming languages. Fourth, (additional) metadata is stored as key-value pairs in plain-text JSON files. A small minority of data in AOMIC is stored using different file formats (such as *hdf5* for composite transforms of MRI data and some Freesurfer files), but these are unlikely to be relevant for most users.

Apart from data *formats*, we can distinguish different data *types* within AOMIC. Following BIDS convention, data types are distinguished based on an “identifier” at the end of the file name (before the extension). For example, T1-weighted files (e.g., *sub-0001_T1w.nii.gz*) are distinguished by the *_T1w* identifier and event log files for task-based functional MRI data (e.g., *sub-001_task-workingmemory_acq-seq_events.tsv*) are distinguished by the *_events* identifier. All data types and associated identifiers within AOMIC are listed in Table 7.

**Table 7.** All data types with associated identifiers, descriptions, and modalities.

| **Raw data** |  |  |
| --- | --- | --- |
| **Identifier (ext.)** | **Description** | **Modality** |
| *_T1w* (nii.gz) | T1-weighted scan | Anatomical MRI |
| *_bold* (nii.gz) | Functional (BOLD) MRI scan | Functional MRI |
| *_magnitude1* (nii.gz) | Average magnitude image (across two echoes) of B0 fieldmap scan (PIOP2 only) | Fieldmap |
| *_phasediff* (nii.gz) | Difference between two phase images of B0 fieldmap scan | Fieldmap |
| *_dwi* (.nii.gz) | Diffusion-weighted scan | DWI |
| *_dwi* (.bvec) | Diffusion gradient table (plain-text file) | DWI |
| *_dwi* (.bval) | Diffusion b-value sequence (plain-text file) | DWI |
| *_physio* (tsv.gz) | Physiology (cardiac/respiratory traces + volume onsets) | Physiology |
| *_events* (tsv) | Onsets, durations, and other relevant properties of events during task fMRI | Behavior |

<sup>3</sup> https://cran.r-project.org/web/packages/gifti/

| * (.json) | Metadata associated with a specific file, acquisition type, or file type | All modalities |
| --- | --- | --- |
| **Derivatives** |  |  |
| *Fmriprep derivatives* |  |  |
| Identifier (ext.) | Description | Modality |
| _T1w (nii.gz) | Preprocessed T1-weighted scan | Functional MRI |
| _mask (nii.gz) | Binary brain mask (0: not brain, 1: brain) | Functional/<br>anatomical/<br>diffusion MRI |
| _dseg (nii.gz) | Discrete segmentation (0: background, 1: gray matter, 2: white matter, 3: cerebrospinal fluid) | Anatomical MRI |
| _probseg (nii.gz) | Probabilistic segmentation (1<sup>st</sup> volume: gray matter, 2<sup>nd</sup> volume: white matter) |  |
| .surf (.gii) | Reconstructed surfaces (e.g., inflated, midthickness, pial, and smoothwm) | Anatomical MRI |
| _xfm (.h5) | Linear + non-linear transformation parameters between two spaces (orig, T1w, MNI152NLin2009cAsym, and fsaverage5) | Functional/<br>anatomical MRI |
| _xfm (.txt) | Linear transformation parameters between two spaces (orig, T1w, MNI152NLin2009cAsym, and fsaverage5) | Functional/<br>anatomical MRI |
| _bold (nii.gz) | Preprocessed BOLD-MRI scan | Functional MRI |
| _boldref (nii.gz) | Reference volume used for computing transformation parameters | Functional MRI |
| _regressors (tsv) | Table with confound regressors for BOLD-MRI data | Functional MRI |
| * (.json) | Metadata from file with the same name (excluding extension) | All modalities |
| *DWI derivatives* |  |  |
| _dwi (.nii.gz) | Preprocessed diffusion-weighted scan | DWI |
| _dwi (.bvec) | Preprocessed gradient table | DWI |
| _dwi (.bval) | Preprocessed b-value sequence | DWI |
| _mask (.nii.gz) | Binary brain mask (0: not brain, 1: brain) | DWI |
| _diffmodel | Estimated parameters from the diffusion tensor model | DWI |
| _EVECS | Eigenvectors from the diffusion tensor model | DWI |
| _FA | Fractional anisotropy map | DWI |
| *VBM derivatives* |  |  |
| GMvolume | Estimated voxelwise gray matter volume map | Anatomical MRI |
| *Freesurfer morphological statistics derivatives* |  |  |
| _stats | Morphological statistics per anatomical region from a particular atlas | Anatomical MRI |
| *Physiology derivatives* |  |  |
| _regressors | Physiology-derived RETROICOR, HRV, and RVT regressors for BOLD-MRI data | Physiology |

#### Data repositories used

Data from AOMIC can be subdivided into two broad categories. The first category encompasses all *subject-level* data, both raw data and derivatives. The second category encompasses *group-level* aggregates of data, such as an average (across subjects) TSNR map or group-level task fMRI activation maps. Data from these two categories are stored in separate, dedicated repositories: subject-level data is stored on OpenNeuro (https://openneuro.org)^87^ and the subject-aggregated data is stored on Neurovault (https://neurovault.org)^88^. Data from each dataset — PIOP1, PIOP2, and ID1000 — are stored in separate repositories on OpenNeuro and Neurovault. URLs to these repositories for all datasets can be found in Table 8. Apart from the option to download data using a web browser, we provide code to download the data programmatically (see https://nilab-uva.github.io/AOMIC.github.io).

**Table 8.** Data repositories for subject data (Openneuro) and group-level data (Neurovault).

|  | **ID1000** | **PIOP1** | **PIOP2** |
| --- | --- | --- | --- |
| Openneuro URL | https://openneuro.org/datasets/ds003097 | https://openneuro.org/datasets/ds002785 | https://openneuro.org/datasets/ds002790 |
| Neurovault URL | https://neurovault.org/collections/7105/ | https://neurovault.org/collections/7103/ | https://neurovault.org/collections/7104/ |

#### Data anonymization

In curating this collection, we took several steps in ensuring the anonymity of participants. All measures were discussed with the data protection officer of the University of Amsterdam and the data steward of the department of psychology, who deemed the anonymized data to be in accordance with the European General Data Protection Regulation (GDPR).

First, all personally identifiable information (such as subjects’ name, date of birth, and contact information) in all datasets were irreversibly destroyed. Second, using the *pydeface* software package, we removed facial characteristics (mouth and nose) from all anatomical scans, i.e., the T1-weighted anatomical scans and (in PIOP2) magnitude and phase-difference images from the B0 fieldmap. The resulting defaced images were checked visually to confirm that the defacing succeeded. Third, the data files were checked for timestamps and removed when present. Lastly, we randomized the subject identifiers (*sub-xxxx*) for all files. In case participants might have remembered their subject number, they will not be able to look up their own data within our collection.

### Technical Validation

In this section, we describe the measures taken for quality control of the data. This is described per data type (e.g., anatomical T1-weighted images, DWI images, physiology, etc.), rather than per dataset, as the procedure for quality control per data type was largely identical across the datasets. Importantly, we take a conservative approach towards exclusion of data, i.e., we generally did not exclude data unless (1) it was corrupted by scanner-related incorrigible artifacts, such as reconstruction errors, (2) when preprocessing fails due to insufficient data quality (e.g., in case of strong spatial inhomogeneity of structural T1-weighted scans, preventing accurate segmentation), (3) an absence of a usable T1-weighted scan (which is necessary for most preprocessing pipelines), or (4) incidental findings. This way, the data from AOMIC can also be used to evaluate artifact-correction methods and other preprocessing techniques aimed to post-hoc improve data quality and, importantly, this places the responsibility for inclusion and exclusion of data in the hands of the users of the datasets.

Researchers not interested in using AOMIC data for artifact-correction or preprocessing techniques may still want to exclude data that do not meet their quality standards. As such, we include, for each modality (T1-weighted, BOLD, and DWI) separately, a file with several quality control metrics across subjects. The quality control metrics for the T1-weighted and functional (BOLD) MRI scans were computed by the Mriqc package and are stored in the *group_T1w.tsv* and *group_T1w.tsv* files in the *mriqc* derivative folder. The quality control metrics for the DWI scans were derived from the output of FSL’s *eddy* algorithm and are stored in the *group_dwi.tsv* file in the *dwipreproc* derivative folder. Using these precomputed quality control metrics, researchers can decide which data to include based on their own quality criteria.

#### T1-weighted scans

All T1-weighted scans were run through the Mriqc pipeline, which outputs several quality control metrics as well as a report with visualizations of different aspects of the data. All individual subject reports were visually checked for artifacts including reconstruction errors, failure of defacing, normalization issues, and segmentation issues (and the corresponding data excluded when appropriate). In Figure 4, we visualize several quality control metrics related to the T1-weighted scans across all three datasets. In general, data quality appears to increase over time (with ID1000 being the oldest dataset, followed by PIOP1 and PIOP2), presumably due to improvements in hardware (see Scanner details and general scanning protocol). All quality control metrics related to the T1-weighted scans, including those visualized in Figure 4, are stored in the *group_T1w.tsv* file in the *mriqc* derivatives folder.

**Figure 4.**
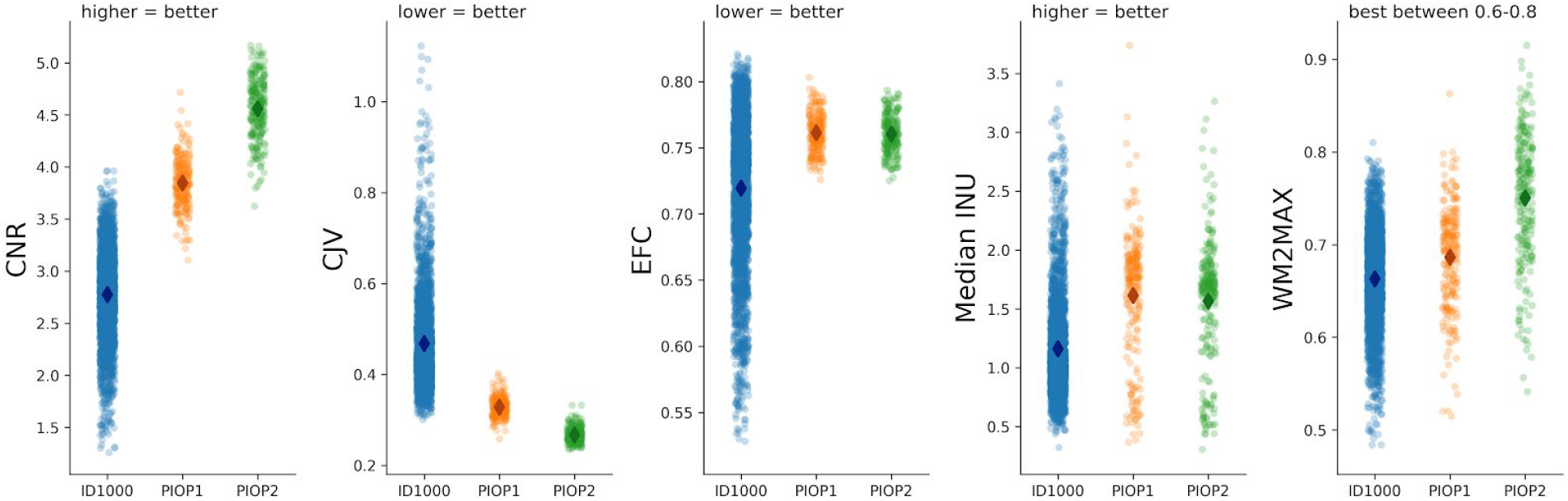
Quality control metrics related to the T1-weighted scans. CNR: contrast-to-noise ratio^89^; CJV: coefficient of joint variation^90^, an index reflecting head motion and spatial inhomogeneity; EFC: entropy-focused criterion^91^, an index reflecting head motion and ghosting; INU: intensity non-uniformity, an index of spatial inhomogeneity; WM2MAX: ratio of median white-matter intensity to the 95% percentile of all signal intensities; low values may lead to problems with tissue segmentation.

#### Functional (BOLD) scans

Like the T1-weighted images, the functional (BOLD) scans were run through the Mriqc pipeline. The resulting reports were visually checked for artifacts including reconstruction errors, registration issues, and incorrect brain masks.

In Figure 5, we visualize several quality control metrics related to the functional scans across all three datasets. Similar to the T1-weighted quality control metrics, the functional quality control metrics indicate an improvement of quality over time. Also note the clear decrease in temporal signal-to-noise ratio (tSNR) for multiband-accelerated scans (consistent with ref^92^). All quality control metrics related to the functional MRI scans, including those visualized in Figure 5, are stored in the *group_bold.tsv* file in the *mriqc* derivatives folder.

**Figure 5.**
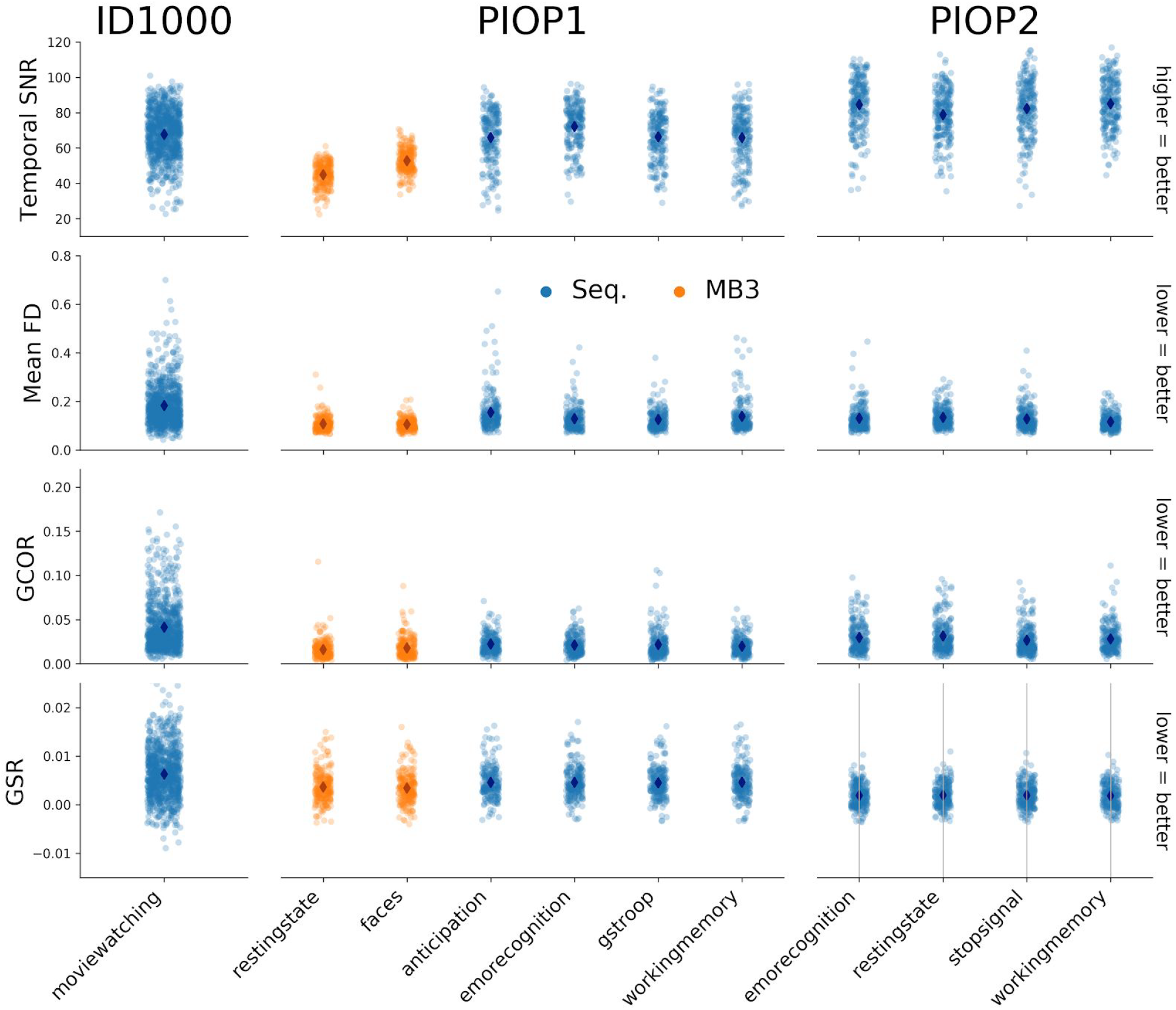
Quality control metrics related to the functional (BOLD) MRI scans. SNR: signal-to-noise ratio, an index of signal quality; FD: framewise displacement^93^, an index of overall movement; GCOR: global correlation^94^, an index of the presence of global signals; GSR: ghost-to-signal ratio, an index of ghosting along the phase-encoding axis.

In addition to global quality control metrics, we also computed whole-brain (voxelwise) temporal signal-to-noise ratio (tSNR) maps, computed by dividing the average signal (across time) by the standard deviation of the signal (across time) of the motion-corrected and spatially normalized data. This is done for each subject and fMRI scan separately, which are subsequently averaged across subjects and scans (but separately for sequential and multiband-accelerated scans in PIOP1 to highlight the tSNR cost of multiband-acceleration).

In Figure 6, we visualize these tSNR maps for each dataset (and separately for the sequential and multiband scans of PIOP1). Again, there appears to be an increase in tSNR across time. Corresponding whole-brain tSNR maps can be viewed and downloaded from Neurovault (i.e., files with the *_tsnr* identifier).

**Figure 6.**
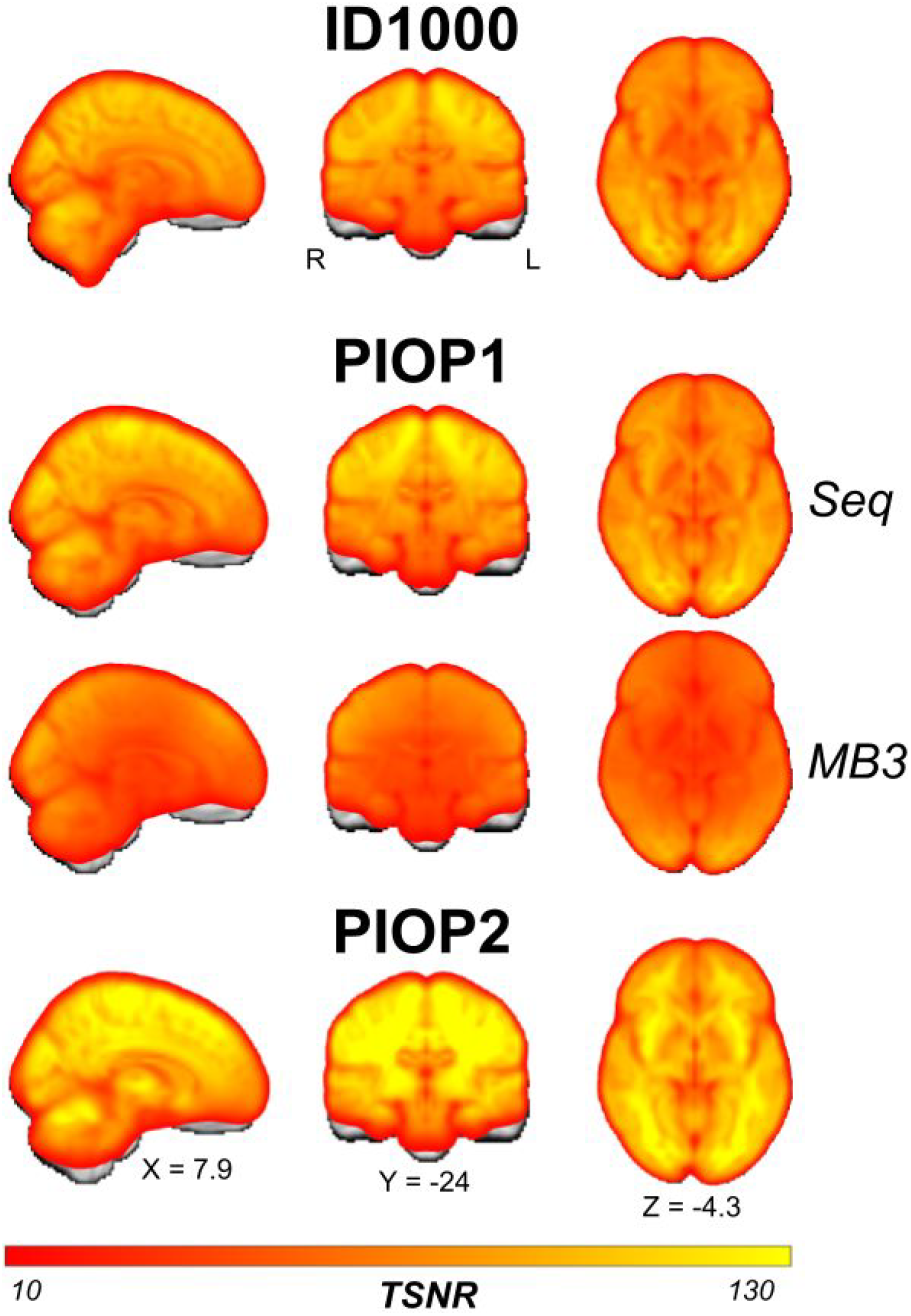
Average (across subjects and runs) temporal signal-to-noise (tSNR) maps of each type of functional (BOLD) MRI scan in each dataset. Unthresholded whole-brain tSNR maps are available on Neurovault.

For the fMRI data with an explicit task (i.e., all fMRI data except for the PIOP resting-state fMRI scans and the ID1000 movie watching fMRI scan), we additionally computed group-level whole-brain statistics maps. To do so, we ran simple first-level models (including task-regressors as well as a discrete cosine basis set functioning as a high-pass filter of 128 seconds and six motion regressors) and computed first-level contrast maps for each subject which were subsequently analyzed in a random effects group-level (intercept) model. In Figure 7, we show the (uncorrected) whole-brain group-level results for each task. Note that we chose these specific contrasts to demonstrate that the tasks elicit to-be expected effects (e.g., amygdala activity in the emotion matching task and cingulate cortex activity in the gender-stroop task). Different, and more sophisticated analyses, including analysis of between-subject factors, are possible with this data and the associated event files.

**Figure 7.**
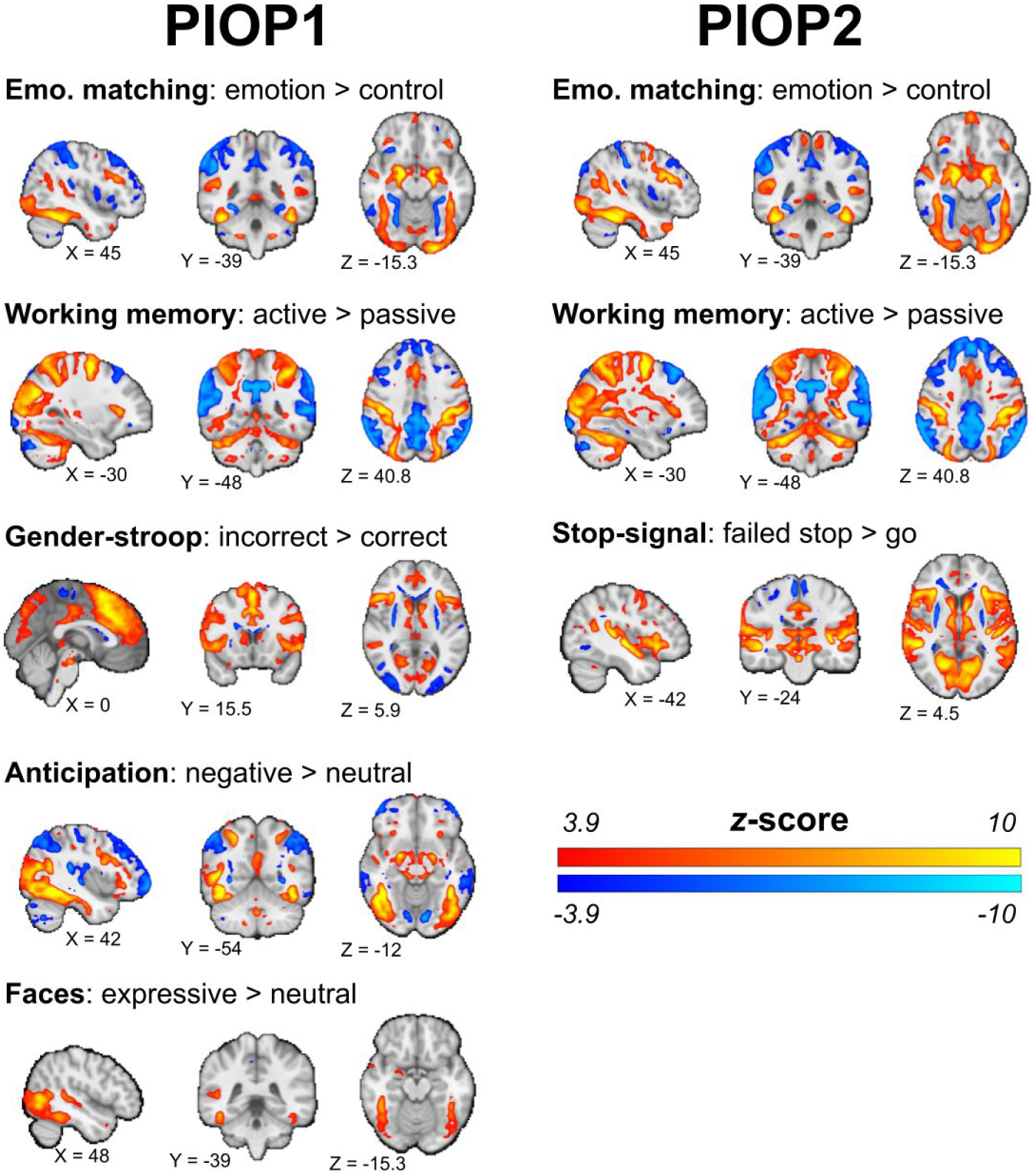
Results from task-specific group-level analyses. Brain maps show uncorrected effects (*p* < 0.00001, two-sided) and were linearly interpolated for visualization in FSLeyes. Unthresholded whole-brain *z*-value maps are available on Neurovault. Unthresholded whole-brain z-value maps are available on Neurovault.

To validate the quality of the resting-state functional MRI scans in PIOP1 and PIOP2, we ran dual regression analyses^95^ using the spatial ICA maps from Smith and colleagues (10-component version)^96^. Figure 8 shows the group-level dual regression results from both PIOP1 and PIOP2 for the first four components next to the original ICA map from Smith and colleagues^96^.

**Figure 8.**
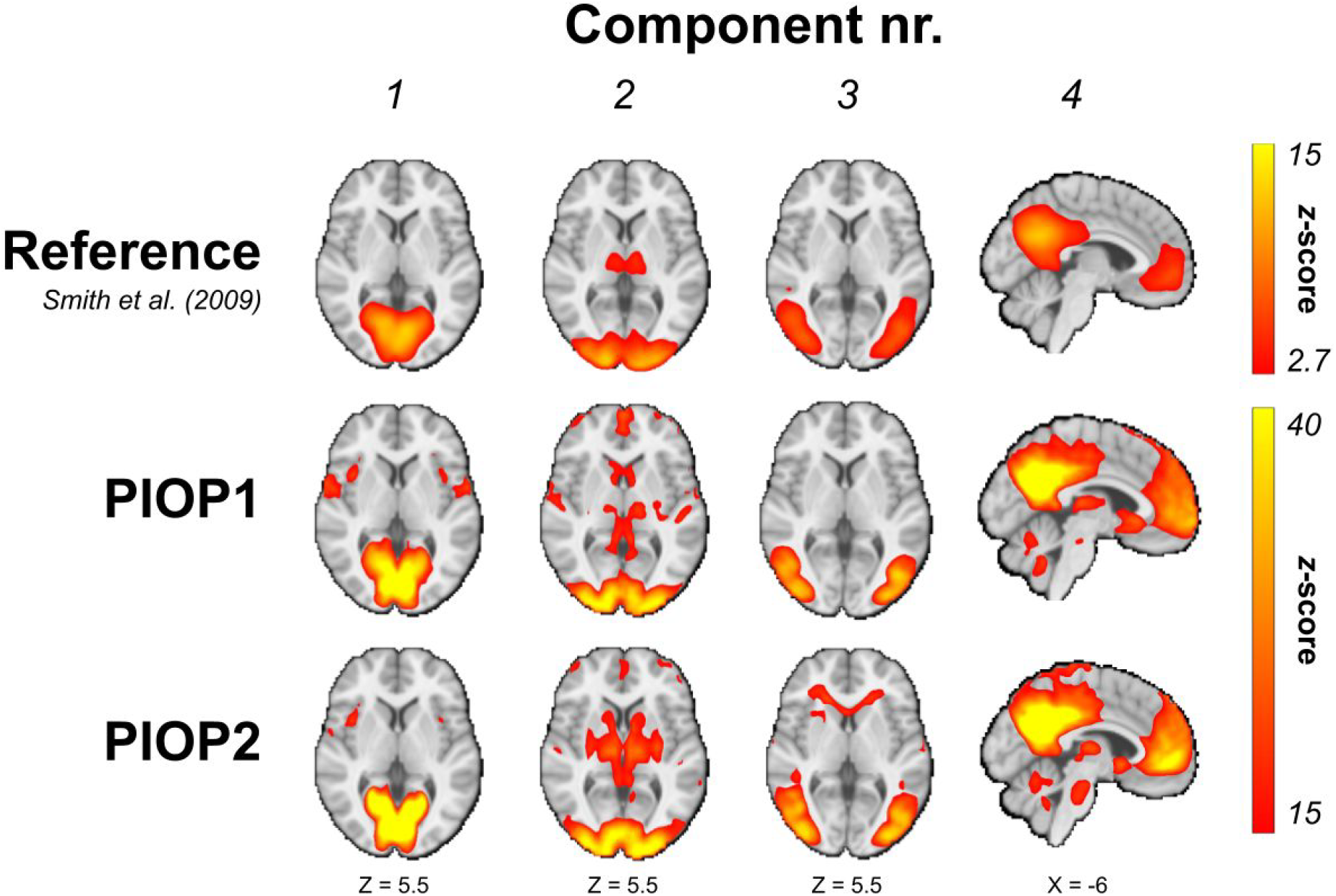
Group-level dual regression results for the first four components of Smith and colleagues^96^. Unthresholded *z*-value maps are available on Neurovault. Unthresholded whole-brain z-value maps are available on Neurovault.

Finally, to assess the quality of the ID1000 functional MRI data, we performed a voxelwise whole-brain “inter-subject correlation” (ISC) analysis, using the BrainIAK software package^83,97^ on data from a subset of 100 participants (randomly drawn from the ID1000 dataset). Before computing the inter-subject correlations, the data were masked by an intersected functional brain mask and grey matter mask (probability > 0.1). Low-frequency drift (with a cutoff of 128 seconds), the mean signal within the cerebrospinal fluid, global (whole-brain average) signal, and six motion parameters were regressed out before computing the ISCs. The average (across subjects) voxelwise ISCs are visualized in Figure 9, which shows the expected inter-subject synchrony in the ventral and dorsal visual stream while watching the video.

**Figure 9.**
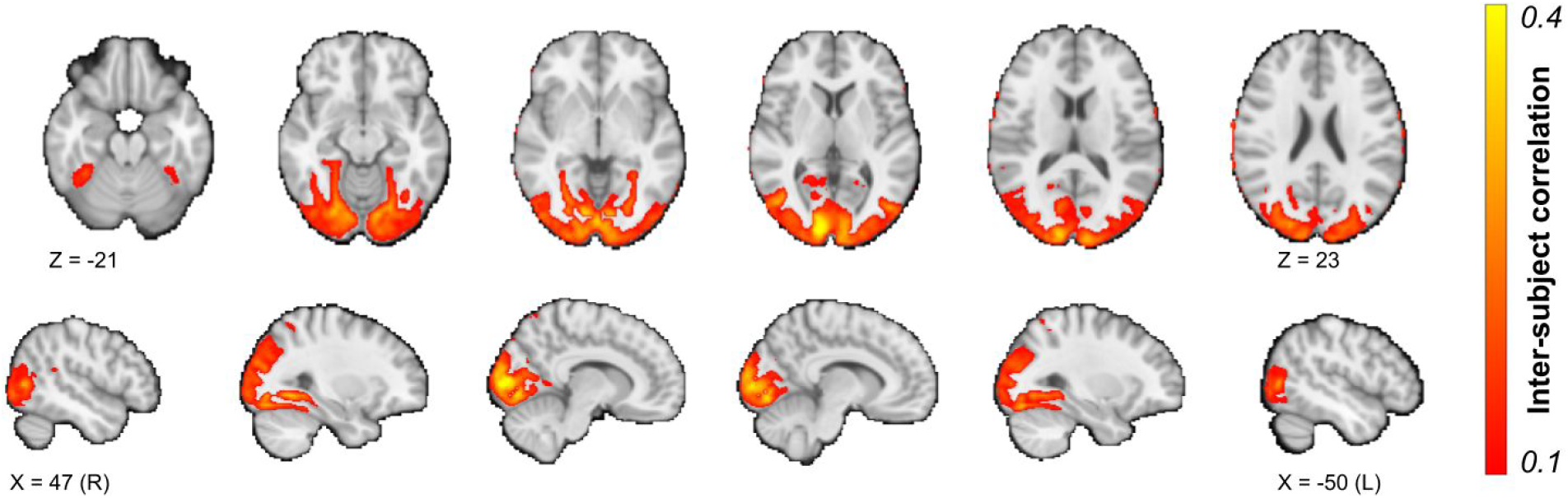
Results from the voxelwise ISC analysis, arbitrarily thresholded at 0.1. An unthresholded whole-brain ISC map is available on Neurovault (https://neurovault.org/images/377226/).

#### Diffusion-weighted scans

Before preprocessing, the b=0 volume from each DWI scan was extracted and visually checked for severe artifacts and reconstruction errors (in which case the data was excluded). After preprocessing and DTI model fitting, we furthermore visualized each estimated fractional anisotropy (FA) map and the color-coded FA-modulated (absolute) eigenvectors for issues with the gradient directions.

Furthermore, we extracted quality control metrics based on outputs from the eddy correction/motion correction procedure in the DWI preprocessing pipeline as implemented in FSL’s *eddy* algorithm (based on the procedure outlined in ref^98^). Specifically, we computed the mean framewise displacement across volumes based on the realignment parameters from motion correction, the percentage of “outlier slices” (as determined by FSL *eddy*) in total and per volume, and the standard deviation of the estimated linear eddy current distortions across volumes. These metrics are visualized in Figure 10. Note that the y-axis for the standard deviation of the eddy currents for ID1000 has a larger range than for PIOP1 and PIOP2 to show the scans with particularly strong eddy current fluctuations.

**Figure 10.**
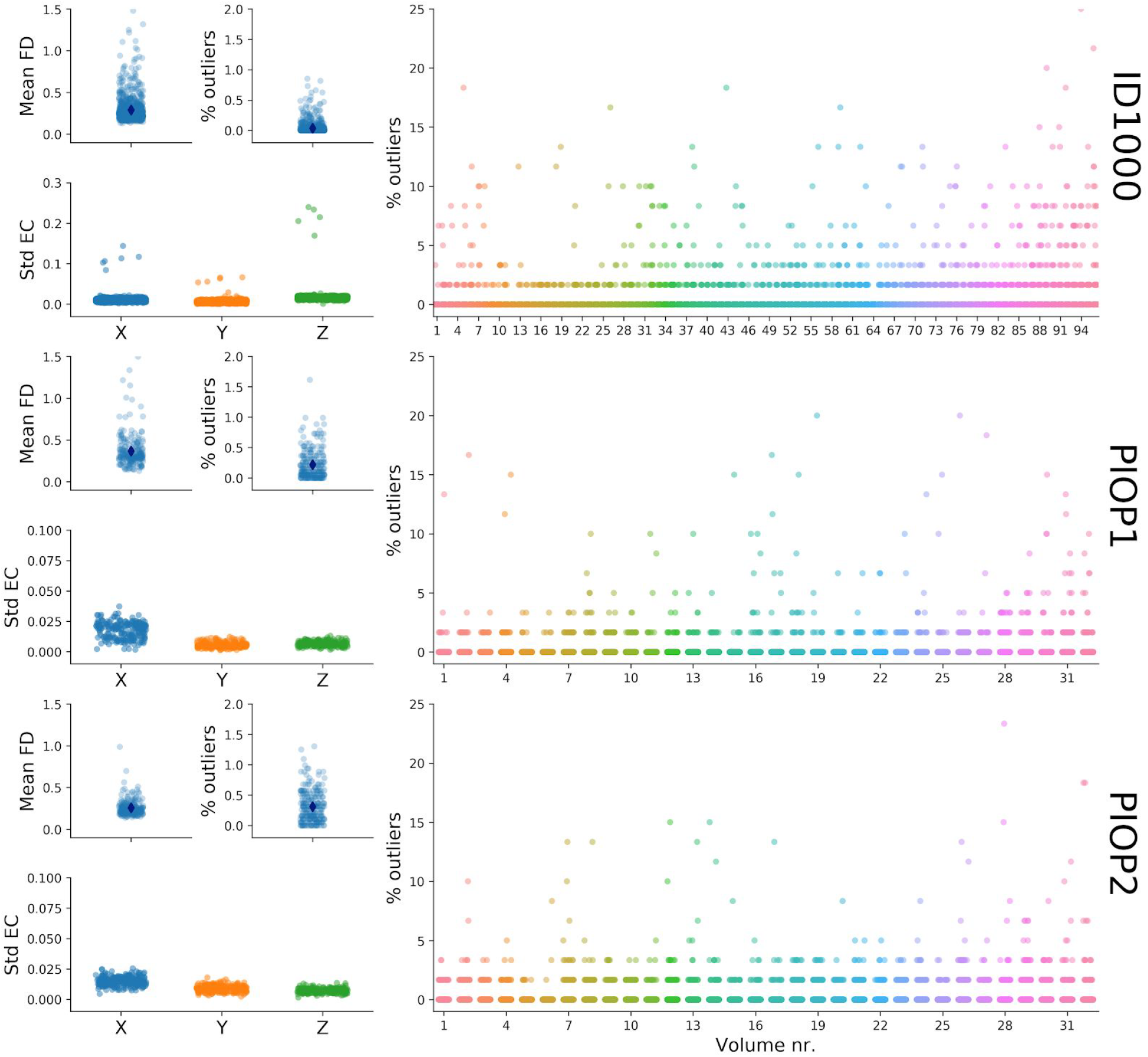
Quality control metrics related to the diffusion-weighted scans. FD: framewise displacement, Std EC: standard deviation of the linear terms of the eddy current distortions in Hz/mm.

Finally, for each dataset, we transformed all preprocessed DTI eigenvectors to a population template estimated on all FA images and computed the voxelwise median across subjects. The median eigenvector images are visualized in Figure 11 as “diffusion-encoded color” (DEC) images, in which values are modulated by the associated FA values.

**Figure 11.**
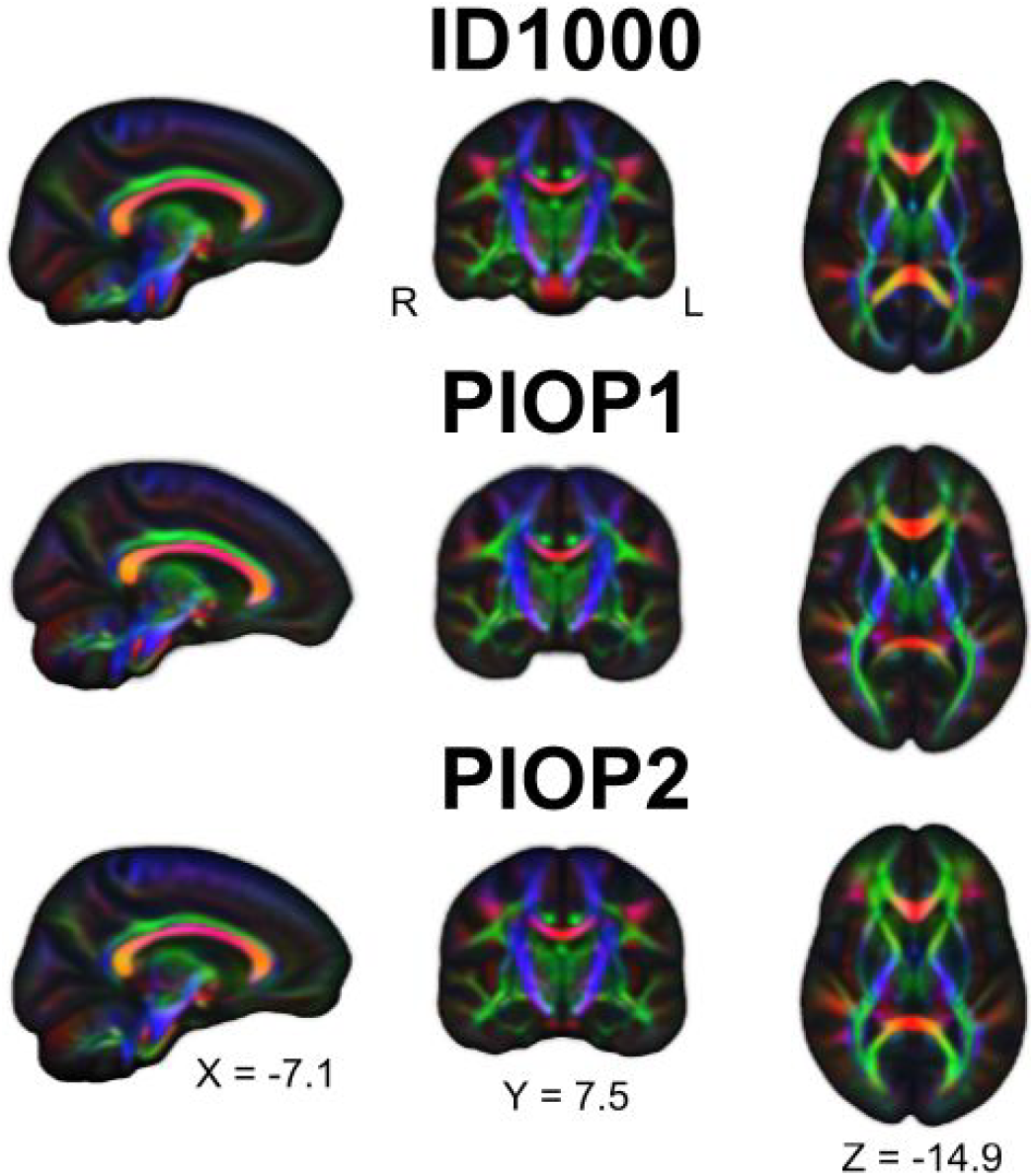
Diffusion-encoded color images of the FA-modulated median DTI eigenvectors across subjects. Red colors denote preferential diffusion along the sagittal axis (left-right), green colors denote preferential diffusion along the coronal axis (anterior-posterior), and blue colors denote preferential diffusion along the axial axis (inferior-superior). Brighter colors denote stronger preferential diffusion.

#### Physiological data

After conversion to BIDS, physiological data was visually checked for quality by plotting the scanner triggers (i.e., volume onsets) and the cardiac and respiratory traces. Files missing a substantial window of data (>10 seconds) were excluded as well as files for which the scanner triggers could not be estimated reliably. Figures of the physiology traces and scanner triggers for each file are included in the physiology derivatives. Additionally, we fit first-level (subject-specific) and subsequently group-level (subject-average) models using the physiology regressors for each dataset. In Figure 12, we visualize the effects of the different RETROICOR components (respiratory, cardiac, and interaction regressors; an *F*-test) and the HRV and RVT regressors (a *t*-test). Unthresholded whole-brain maps are available from Neurovault.

**Figure 12.**
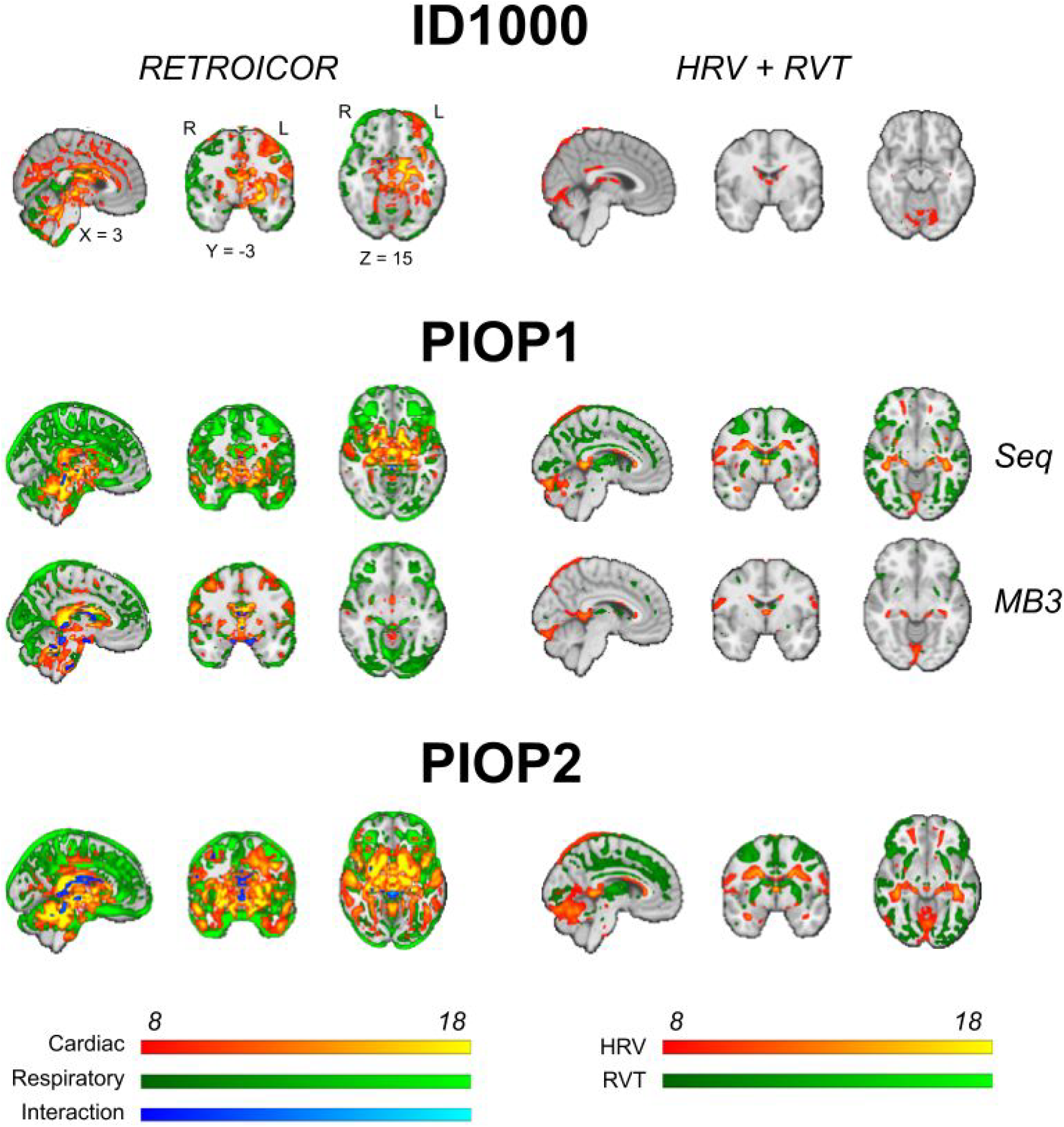
Results from group-level physiology analyses. Brain maps show uncorrected effects (thresholded arbitrarily at *z* > 8) and were linearly interpolated for visualization in FSLeyes. Unthresholded whole-brain z-value maps are available on Neurovault.

#### Psychometric data

The patterns of correlations within the scales of the questionnaires are consistent with those reported in literature, indicating that this data is overall reliable. The pattern of correlations between scales of different questionnaires and external variables is also consistent with those reported in literature and what would be expected on theoretical grounds.

##### Intelligence Structure Test (IST)

The subscales of the IST (fluid and crystallized intelligence and memory) are strongly correlated with each other. The validity of the measure data is supported by the correlation with relevant external variables like educational level, *r*(926) = 0.46) and background SES, *r*(926) = 0.35 (see Table 9).

**Table 9.**
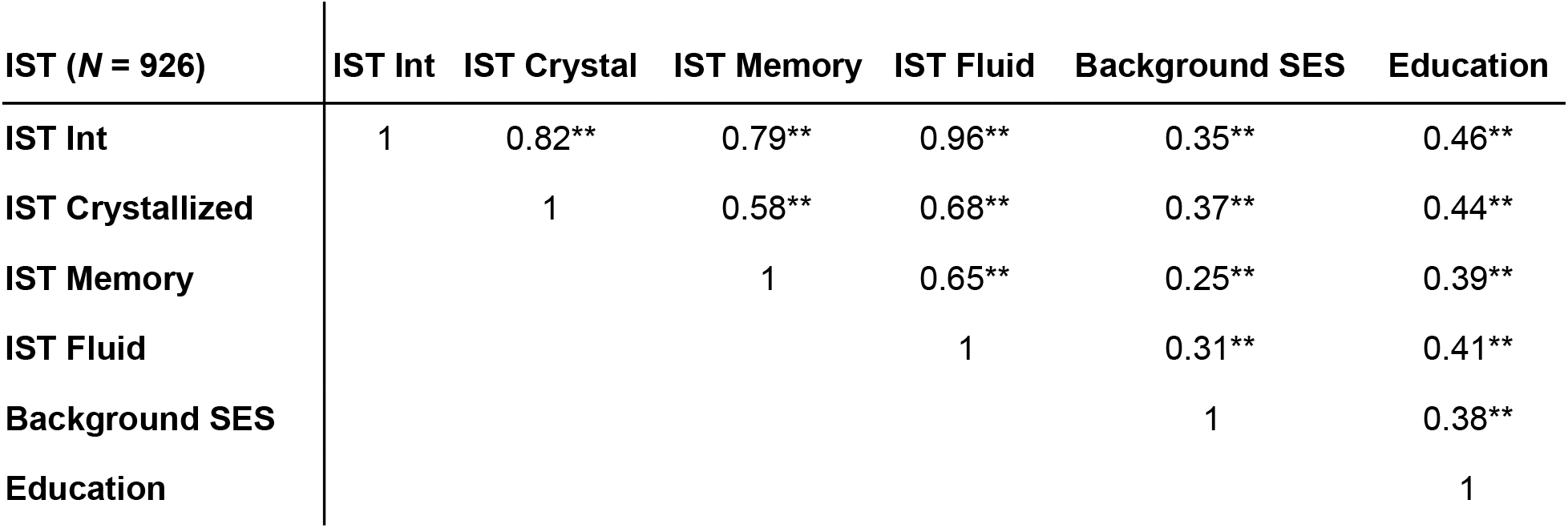
Correlations between total score, and subscales of the IST and relevant external variables (background SES and educational level). IST: Intelligence Structure Test, Int: Total Intelligence, SES: background social-economic status. ** indicates *p* < 0.01.

| IST (N = 926) | IST Int | IST Crystall | IST Memory | IST Fluid | Background SES | Education |
| --- | --- | --- | --- | --- | --- | --- |
| IST Int | 1 | 0.82** | 0.79** | 0.96** | 0.35** | 0.46** |
| IST Crystallized |  | 1 | 0.58** | 0.68** | 0.37** | 0.44** |
| IST Memory |  |  | 1 | 0.65** | 0.25** | 0.39** |
| IST Fluid |  |  |  | 1 | 0.31** | 0.41** |
| Background SES |  |  |  |  | 1 | 0.38** |
| Education |  |  |  |  |  | 1 |

##### Personality: NEO-FFI

The cross-correlation patterns of the 5 NEO-FFI scales are depicted in table 10. Significant correlations exist between the scales, and the correlation pattern is overall consistent with the reported norm data for this test^18^. The correlation between cross-correlation patterns of the three datasets is very consistent (*r* = 0.88 between PIOP1 and PIOP2, and on average *r* = 0.74 between ID1000 and PIOP), with as a notable outlier a negative correlation, *r*(928) = −0.13, *p* < 0.001, between extraversion and agreeableness in the ID1000 dataset and a positive correlation for these scales in the PIOP1, *r*(216) = 0.20, *p* < 0.005, and PIOP2, *r*(226) = 0.26, *p* < 0.001. A source for this discrepancy could be the difference in population sample between the PIOP1 and PIOP2 studies and the ID1000 study.

**Table 10.**
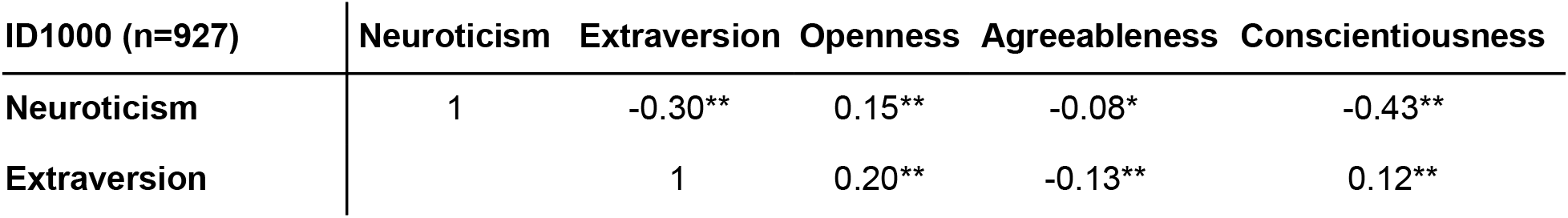
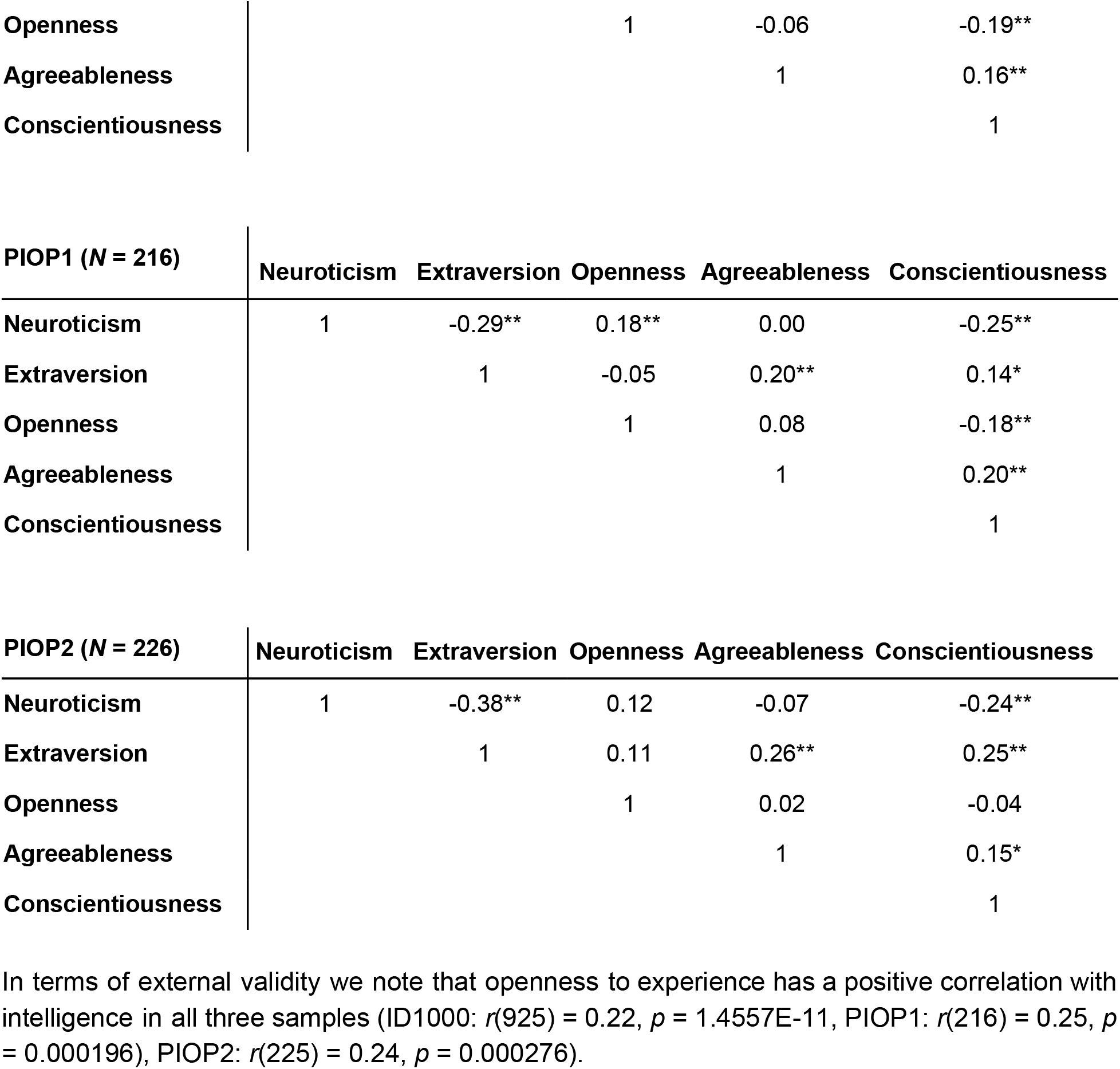
Cross-correlations for the subscales of the NEO-FFI for the ID1000, PIOP1 and PIOP2 samples. With the exception of the correlation between agreeableness and extraversion the cross-correlation patterns are very similar across samples. * indicates p<0.05. ** indicates p<0.01.

In terms of external validity we note that openness to experience has a positive correlation with intelligence in all three samples (ID1000: *r*(925) = 0.22, *p* = 1.4557E-11, PIOP1: *r*(216) = 0.25, *p* = 0.000196), PIOP2: *r*(225) = 0.24, *p* = 0.000276).

##### BIS/BAS

The cross-correlation patterns of the BIS/BAS scales are depicted in table 11. The cross-correlation between the scales are similar to the one reported by Franken and colleagues^16^ and contrary to what Carver & White^34^ predicted, with a positive correlation between the three different BAS-scales, but also between BIS and BAS-Reward, *r*(927) = 0.194.

**Table 11.**
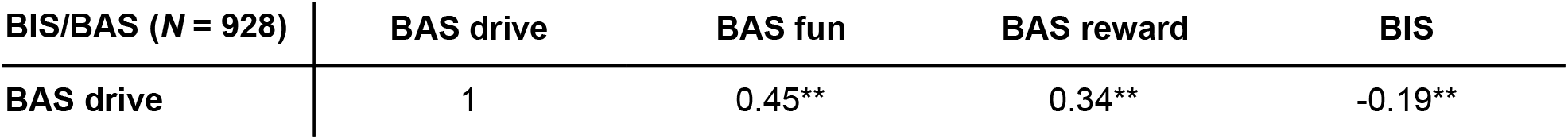
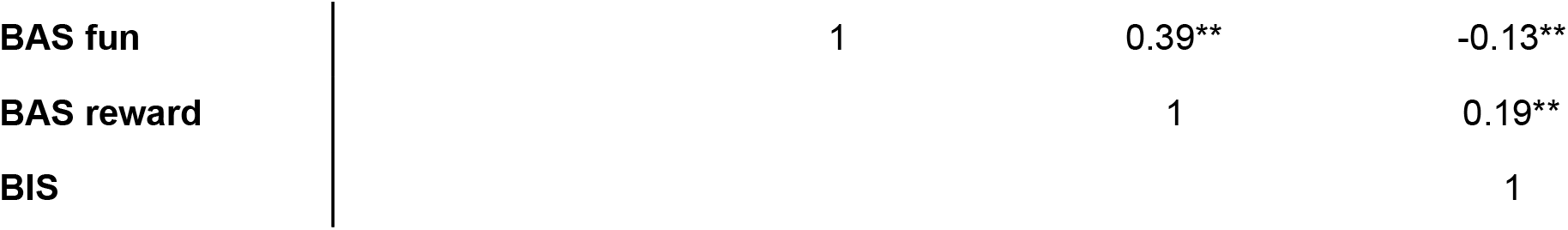
Cross-correlations for the subscales of the BISBAS for the ID1000 sample. The pattern of correlations is consistent with that reported in literature.

The STAI-T scale measures trait anxiety. Because this instrument only consists of one scale we evaluate its reliability on the degree in which it shows correlations with other questionnaire scales that also have a pretension of measuring negative emotionality. Because we observe positive correlations with both Neuroticisms, *r*(927) = 0.77, *p* = 1E-183) and BIS, *r*(927) = 0.514, *p* = 1E-63) we conclude that the reported scales are reliable and consistent.

## Acknowledgements

We thank all research assistants and students who helped collecting the data of the three projects, Jasper Wijnen and Marco Teunisse for advice and guidance with respect to anonymization and GDPR-related concerns, and Jos Bloemers, Sennay Ghebreab, Adriaan Tuiten, Joram van Driel, Christian Olivers, Ilja Sligte, Sara Jahfari, Guido van Wingen, and Suzanne Oosterwijk for help with designing the paradigms, Marcus Spaan for technical support, and Franklin Feingold and Joe Wexler for help with uploading the datasets to Openneuro. The acquisition of the ID1000 dataset was funded by a grant from the Alan Turing Institute, Almere, The Netherlands.

## Author contributions

LS curated the datasets, did visual quality control, preprocessed and analyzed the data, and wrote the article.

MMvdM did visual quality control, analyzed the data, and wrote the article.

TB designed the data acquisition protocol, collected the data, and did visual quality control for PIOP1 and PIOP2.

AvdL designed experiments, collected data, and did visual quality control for ID1000.

AE designed and implemented the questionnaires in ID1000, PIOP1 and PIOP2 and managed questionnaire data from ID1000.

HSS was principal investigator of the ID1000, PIOP1, and PIOP2 studies, designed the data acquisition protocol of ID1000, and wrote the article.

## Footnotes

#1 https://nipy.org/nibabel/

#2 https://cran.r-project.org/web/packages/oro.nifti

#3 https://cran.r-project.org/web/packages/gifti

